# Strength of developmental carry-over varies continuously with environmental conditions in a wild bird population

**DOI:** 10.64898/2026.09.18.752706

**Authors:** Devi Satarkar, David López-Idiáquez, Louis Bliard, Irem Sepil, Ben C. Sheldon

## Abstract

Early-life phenotypes frequently predict adult phenotypes through developmental ‘silver-spoon’ carry-over effects, but how strongly and under what circumstances? Using 40 years of individual-based data from a great tit (*Parus major*) population, we used covariance reaction norms to understand how the natal-to-adult body mass association shifts across temperature, rainfall, and population density gradients during both life stages. We show that the natal–adult mass correlation is substantial (r ≈ 0.7) but strongly environment-dependent. Reaction norm shapes across climatic and density axes point to distinct mechanisms, including compensatory growth under benign adult conditions and density-dependent modulation of early-life differences. We further show that the conditions experienced in adulthood largely drive these patterns and determine whether early-life advantages are retained, even when explicitly tested within an environmental-matching paradigm. Decomposing the correlation using 38 generations of pedigree data revealed no evidence of gene-by-environment interactions. Instead, the observed phenotypic correlation reaction norms could be attributed almost entirely to environmentally-sensitive non-genetic individual differences. Overall, our findings show that climatic and demographic shifts can substantially reshape developmental carry-over without altering the genetic architecture tying body mass across life stages, with implications for how reliably early-life mass can predict individual quality and adult traits in wild animal populations.

## Introduction

The environmental conditions an individual experiences during development can have lasting consequences for phenotype and fitness well into adulthood, a pattern documented across many taxa, including birds and mammals (Lea & Rosebaum, 2020; Lindström, 1999; Monaghan, 2007). Individuals that develop under more favourable conditions tend to grow faster, attain greater body mass, survive better, and achieve higher reproductive success than those whose development was constrained (Lindström, 1999; Pigeon et al., 2019; Reid et al., 2003). Grafen (1988) referred to this phenomenon as the ‘silver-spoon’ effect. However, the linkage between early-life and later-life phenotypes can take many forms and be driven by multiple processes.

This carry-over linkage may vary with the environmental conditions experienced over an individual’s lifetime (Fig. 1). For instance, the adult environment may modulate the strength of carry-over through compensatory mechanisms. When resources are abundant, individuals that developed under poor conditions may have the opportunity to catch up and partially close the gap with higher-quality individuals, weakening the predictive link between early-life and adult body mass. When adult conditions are harsh and resources scarce, early-life differences may persist because lower-quality individuals cannot compensate (Metcalfe & Monaghan, 2001). In such scenarios, the carry-over association would therefore be expected to be weakest in benign adult environments and strongest in harsh ones (Fig. 1D). In contrast, the environmental matching or predictive adaptive response hypothesis (Bateson et al., 2014) suggests that individuals may perform better when adult environments resemble natal ones, regardless of the environmental quality, because organisms are thought to adjust their developmental trajectory in response to early-life environmental cues, effectively calibrating their phenotype to anticipated future conditions (Monaghan, 2007) (Fig. 1E).

**Figure 1.**
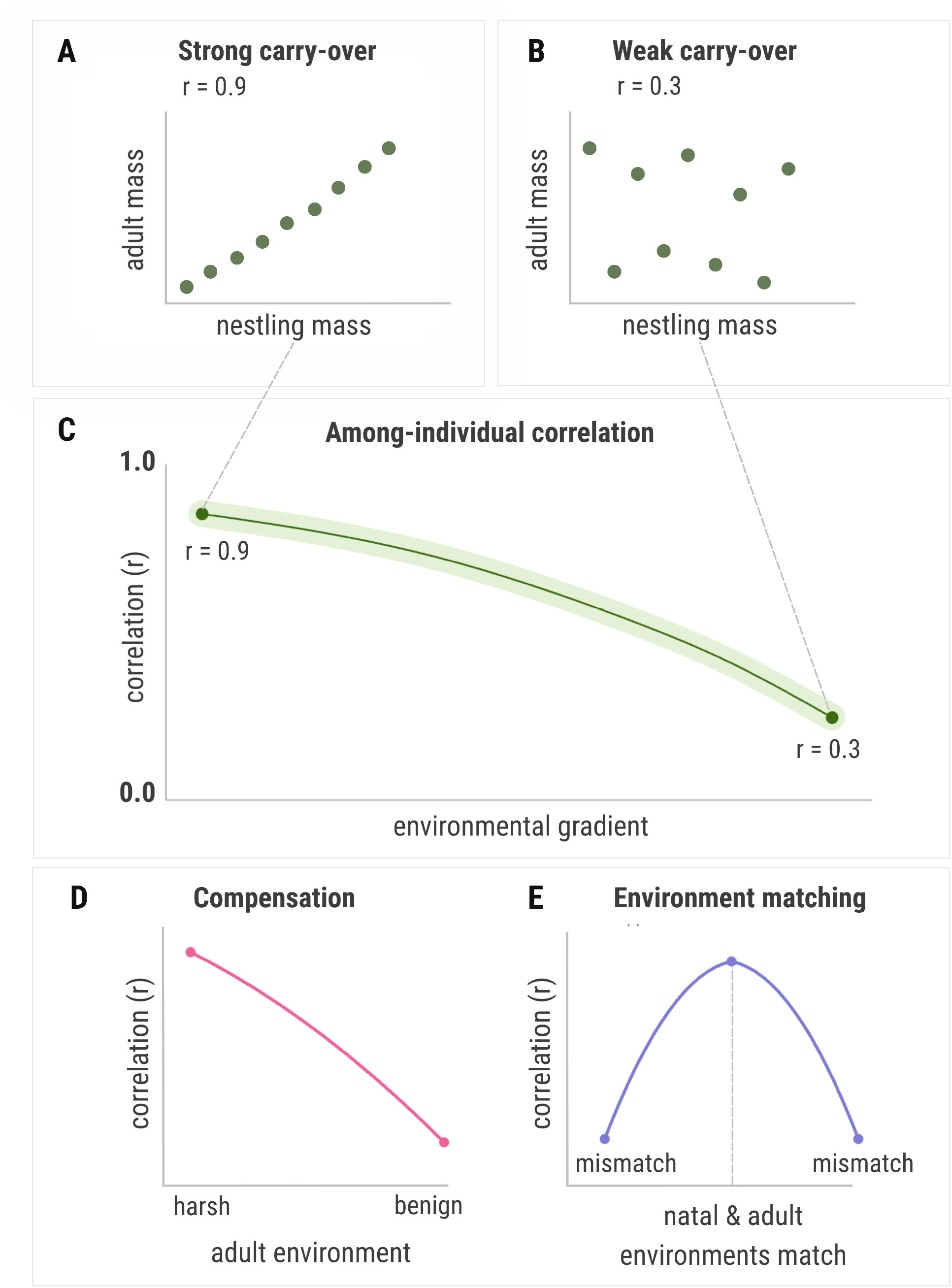
**(A,B)** Illustrative plots of strong versus weak among-individual correlation between nestling mass and adult mass; each point represents an individual and the grey line is a linear fit shown for visual reference. **(C)** A schematic correlation reaction norm translating these two scenarios (A,B) onto a correlation-versus-environment axis; and the format used throughout Figure 2–4, where the y-axis tracks how consistently natal body mass predicts adult mass. Curve shape is illustrative, and does not represent real data. **(D,E)** Predicted correlation reaction norm shapes under the two frameworks outlined in the introduction: Compensation predicts that the correlation weakens as adult conditions become more benign (D), while environmental matching predicts the correlation is strongest when natal and adult environments are similar and weakens as they diverge in either direction, regardless of environmental quality (E).

These two frameworks are not mutually exclusive and may operate through distinct mechanisms across different environmental axes (Monaghan, 2007; Pigeon et al., 2019). For example, temperature may influence developmental calibration through direct physiological effects on growth (Andreasson et al., 2018), whereas variation in resource availability may act more through compensatory pathways (Metcalfe & Monaghan, 2001). Testing these ideas, therefore, requires quantifying carry-over strength as a continuous function of multiple environmental axes. However, most studies examining context-dependent carry-over effects have done so along a single environmental axis in isolation, or by comparing carry-over across discrete environmental categories (Beltran & Tarwater, 2024), such as good versus bad years or high versus low density, rather than estimating it as a continuous function of the environment (Both et al., 1999; Clutton-Brock et al., 1987; Mueller et al., 2025; Pigeon et al., 2019). As a result, the extent to which environmental modulation of carry-over reflects compensatory processes, environmental matching, or a combination of the two remains unclear, particularly across more than one environmental dimension.

Body mass is a particularly useful trait for understanding these processes because it reflects the resources available for growth and energy storage during development (Labocha & Hayes, 2012), predicts fitness-related outcomes including metabolic rate (Hudson et al., 2013), survival probability (Bouwhuis et al., 2015; Mueller et al., 2025), and reproductive output (Maldonado-Chaparro et al., 2015), and despite having a moderate genetic basis (Dumas et al., 2024; Garant et al., 2004), is an inherently labile trait sensitive to variation in the biotic and abiotic environment (Garant et al., 2004; Satarkar et al., 2026). Tracking body mass across life stages provides a useful empirical window onto carry-over effects at the individual level, especially in longterm studies of wild populations where individuals can be measured repeatedly and their developmental histories traced. Yet, the natal-to-adult association in body mass is typically quantified as a single regression coefficient, a population average that implicitly treats carry-over strength as fixed across individuals, years, and environments (López-Idiáquez, Cole, Satarkar, et al., 2026). Understanding how strongly early-life mass predicts adult mass, and whether that association itself changes with environmental conditions across individuals’ lives, remains poorly understood, despite its implications for how individual quality is maintained across life stages.

A further challenge is that the population-level correlation between natal and adult body mass conflates two biologically distinct sources of resemblance: shared genetic effects (including additive, dominance, and epistatic components) and the persistence of non-genetic individual differences across life stages, such as territory quality, parental effects, or early environmental advantage (Dingemanse & Dochtermann, 2013; Kruuk, 2004). These two components have different evolutionary implications. Additive genetic covariance between natal and adult body mass determines how selection on early-life mass is reflected in the adult phenotype across generations, whereas the non-genetic component reflects the degree to which early environmental advantages are maintained or eroded by adult conditions. Whether these components respond differently to the environment, and whether the environmental modulation of carry-over strength is genetic in origin, non-genetic, or both, has direct consequences for predicting how populations may respond to ongoing environmental change.

The recently developed covariance reaction norm (CRN) framework (Martin, 2025) addresses these gaps in a single modelling approach by estimating among-individual covariance between traits as a continuous function of environmental context. Figure 1 illustrates how among-individual correlations are interpreted within this framework for our study. We applied this method to four decades of individual-based data from the great tit (*Parus major*) population at Wytham Woods, UK, (Perrins, 1965), where nestlings are individually ringed and weighed at a standardised age each spring, and those that survive to breed are recaptured and weighed as breeding adults in subsequent years. This allows the natal and adult body mass of the same individuals to be tracked across life stages and across the full range of environmental variation present over the study period. A social pedigree from this population, spanning 38 generations, further allows the estimation of the genetic component underlying the natal-to-adult body mass association.

We had three aims: first, to quantify how natal and adult temperature, rainfall, and population density jointly and continuously modulate the among-individual correlation between nestling and adult body mass. Second, to test whether the degree of cross-life-stage environmental mismatch in temperature, rainfall, and density modulates the carry-over correlation, and what this implies for the environmental matching framework. Third, to decompose the phenotypic carry-over correlation into its genetic and non-genetic components, and test whether each varies across temperature and density gradients. This is particularly relevant under increasing climate variability (IPCC, 2023), where individuals are more likely to develop and breed in highly variable thermal environments within their lifetimes and the extent to which the genetic covariance underlying carry-over is contingent on these environmental conditions will determine how selection on early-life body mass propagates across generations.

## Methods

For this study, we used the ‘covariance reaction norm’ (CRN) framework recently developed by Martin (2025), to estimate the among-individual correlation between natal and adult body mass as a continuous function of natal and adult environmental conditions. The CRN approach extends multivariate animal models to allow trait covariances to vary in response to environmental predictors, making it possible to test how the predictive link between early and late life body condition changes across environments experienced at different life stages. We applied this method to 40 years of individual-based breeding data from the great tit population at Wytham Woods, following the implementation in Martin (2025) and Bliard et al. (2026).

### Study system and data

Great tits (*Parus major*) have been monitored continuously at Wytham Woods, Oxford, UK (51°46′N, 1°20′W) since 1947 using a nestbox population that enables near-complete recording of breeding events within the study area (Perrins, 1965). Each spring, all occupied nestboxes are visited regularly to record detailed breeding data including laying date, clutch size, hatch date, and brood size. Nestlings are individually ringed and weighed to the nearest 0.1 g on day 15 post-hatching, providing a standard measure of prefledging body condition at a stage when nestling mass typically becomes asymptotic (Bouwhuis et al., 2015). Adults are caught and weighed at the nest during chick rearing when their offspring are between 10 to 15 days old, and are individually ringed if not previously tagged (see López-Idiáquez et al., (2026) for details on how body mass is measured in the Wytham tit population).

Our analysis focused on individuals for which both a natal body mass measurement (nestling mass at day 15) and at least one adult body mass measurement from a subsequent breeding attempt were available, and for which both natal and breeding environments were known in detail. After filtering to complete cases across all predictor variables, the final dataset comprised 6,362 breeding records from 4,345 individuals, with birth cohorts spanning 1983–2023 and breeding seasons spanning 1984–2024 (see Supp. Info S2 for a full description of data coverage, and a robustness check using imputed missing data).

Environmental predictors for our models corresponded to either the natal or the breeding environment. For each individual, we calculated the mean daily temperature and mean daily rainfall separately for two individual-specific relative windows: the natal period (the 15-day window from hatching to weighing), and the adult breeding period (the equivalent 15-day window post-hatching of the focal individual’s own offspring). Using individual-specific relative windows rather than fixed seasonal averages, as used in previous studies in this population (López-Idiáquez, Cole, Regan, et al., 2026; Satarkar et al., 2026), better captures the thermal and precipitation conditions actually experienced by each individual during growth and provisioning, since phenological variation among individuals means that birds hatching at different dates can experience substantially different weather conditions within the same season. Daily temperature and precipitation data were obtained from the Met Office Hadley Centre datasets for central England (https://www.metoffice.gov.uk/hadobs/). Phenological mismatch was an additional natal environmental variable, calculated as the absolute difference between the half-fall date of winter moth (*Operophtera brumata*) larvae, the standard index of timing of peak caterpillar abundance in this population (Morley et al., 2025), and the tenth day post-hatching of the focal individual, when the nutritional demands of the nestlings are highest (López-Idiáquez, Cole, Satarkar, et al., 2026). Population density was calculated as the total number of breeding pairs of great tits and blue tits (*Cyanistes caeruleus*) recorded in the nestbox population each year, providing a measure of intra- and inter-specific competition for resources. We combined both species into a single density measure as they both contribute to the resource competition experienced by breeding great tits, albeit via different mechanisms: great tits primarily through interference competition and blue tits through exploitation of shared food resources (Dhondt, 2023; López-Idiáquez, Cole, Satarkar, et al., 2026).

### Statistical analysis

#### Defining environmental contexts

A key modelling decision concerns how to define environmental ‘contexts’, or the units within which among-individual covariances are estimated, and which corresponds to the defined environment that is shared by a group of individuals. The CRN model requires that each context contains individuals measured for both traits, and that context-level environmental predictors are ecologically meaningful for those individuals. We defined contexts as unique birth year x breeding year combinations, so that each context contained only individuals belonging to the same birth and breeding cohort (Fig. S1). This paired context design ensured that individuals experiencing similar natal and breeding environmental conditions were grouped together. Individuals could appear in multiple contexts if they bred in multiple years, but contributed only a single natal mass record measured in their birth year. In our dataset, there were 113 paired contexts with at least 10 individuals spanning the full study period.

Context-level environmental variables were derived from the individual-level measurements described above. For temperature and rainfall, the context-level value was calculated as the mean of the individual-specific values for all birds present in that context. For example, for a context defined by birth year 2002 and breeding year 2004, natal temperature would be calculated for all birds born in 2002 that went on to breed in 2004, as the mean of mean daily temperatures experienced by each bird from 0-15 days post hatching. The adult temperature would be the mean of individual 0-15 day mean temperatures experienced by those same birds when they raised their own offspring in 2004. This approach means that context-level temperature and rainfall reflect the actual thermal and precipitation conditions experienced by the specific cohort of individuals in that context, rather than a generic seasonal average for that year. Population density, being a year-level variable, was assigned directly: natal density corresponds to the total number of breeding pairs in the birth year of the context, and adult density to the total number of breeding pairs in the breeding year. Collinearity among the six environmental predictors was assessed via pairwise correlations and variance inflation factors (VIF) prior to modelling; all predictors were sufficiently independent (VIF < 3) with the exception of a moderate correlation between natal and adult population density (r = 0.59; Fig. S2), reflecting temporal autocorrelation in population size.

## Model structure

We modelled nestling mass (z_1_) and adult mass (z_2_) jointly as Gaussian traits, following a CRN approach (Bliard et al., 2025) where the two traits have different numbers of observations per individual. Nestling mass was measured once per individual (on day 15 post hatching; N = 4,345 nestlings), while adult mass was measured once per breeding attempt, meaning the same individual may contribute multiple observations across different breeding years (N = 6,362 breeding attempts). Each trait was modelled as;

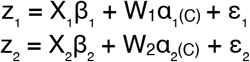

where X_1_ and X_2_ are individual-level matrices of fixed effect predictors including an intercept, W_1_ and W_2_ are matrices that index observations to individual identities across repeated breeding attempts, α_(C)_ are among-individual random effects that vary across environmental contexts C, and ε_1_ and ε_2_ are observation-level residual terms. All response and predictor variables were standardised to zero mean and unit variance prior to analysis.

For nestling mass, fixed effects (X_1_) included natal lay date (i.e., the date the clutch they belonged to was initiated), natal brood size, natal mean temperature, natal mean rainfall, phenological mismatch, and birth year (as a continuous variable), to account for any long-term directional change in nestling mass (López-Idiáquez, Cole, Satarkar, et al., 2026). For adult mass, fixed effects (X_2_) included laying date, mean temperature, mean rainfall, number of offspring, age, sex, and breeding year (as a continuous variable). Fixed effects here serve to account for predictable sources of mean variation in each trait before the covariance structure is estimated, ensuring that the among-individual correlation reflects stable individual differences in condition rather than shared responses to measured environmental variables.

Additionally, birth year and breeding year random effects were included (as categorical variables) for nestling mass and adult mass respectively, to account for unmeasured inter-annual variation in trait means not captured by the fixed predictors. Brood identity and breeding nestbox were also included as random effects, to account for the non-independence of nestlings raised in the same brood and the potential influence of variation in territory quality on adult mass respectively.

The central quantity of interest to be obtained from these analyses is the among-individual covariance between the two traits, specifically, the tendency for individuals who were heavier as nestlings to also be heavier as adults across their breeding lives. After accounting for the above-mentioned fixed effects and random effects, the among-individual effects [α_1(C)_, α_2(C)_] are drawn jointly from a bivariate normal distribution whose covariance matrix P_(C)_ is allowed to vary as a function of environmental context C:

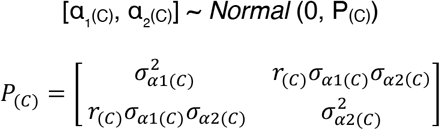

Here 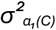 and 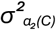 are the among-individual variances in chick weight and adult weight respectively within context C, and *r*_*(C)*_ is the among-individual correlation between them. The among-individual correlation and standard deviations are modelled as linear functions of context-level environmental predictors, contained in matrix X_C_, via appropriate link functions:

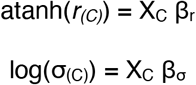

The log link ensures variances remain positive and the inverse hyperbolic tangent function atanh, analogous to a logistic regression but with bounds at −1 and +1 rather than 0 and 1, constrains the estimated correlation to its valid range while allowing additive environmental effects to be estimated on an unbounded scale (Martin, 2025). The intercept of β_r_ gives the baseline among-individual correlation at mean environmental conditions; each slope in β_r_ quantifies how the correlation changes per one standard deviation shift in the corresponding environmental predictor, with all other predictors held at their mean. Slopes of β_r_ indicate how the environment is modulating the carry-over relationship between natal condition and adult body mass.

Due to the nature of available data, we faced constraints regarding the residual terms. Since nestling mass is measured only once per individual, the residual term ε_1_ captures measurement error only, as there is no within-individual variation to separate from measurement noise for this trait. For adult mass, ε_2_ represents genuine within-individual variation across breeding attempts as well as measurement error, though each individual contributed only one adult mass record per context. We therefore faced a non-identifiability issue in the model structure: the baseline among-individual variance (the intercept of log(σ_(C)_)) and the residual standard deviation cannot be estimated simultaneously, because both inflate total phenotypic variance and are indistinguishable when each individual contributes only one mass record per context. Rather than estimating observation-level variance, which aggregates variation due to both among- and within-individual differences, potentially leading to risk of inferential bias (see Figure 1 in Bliard et al., 2025), we fixed the residual standard deviations to externally derived values, allowing us to estimate the among-individual variation (the quantity we were primarily interested in). For nestling mass, we fixed 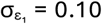, a value broadly consistent with the known precision of the scales used to weigh nestlings, following recommended practice for fixing such terms from external knowledge when repeated measurements are unavailable (Ponzi et al., 2018). For adult mass, we fixed 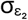 to the REML estimate of within-individual variance from a univariate linear mixed model fitted in *lme4* to the same analytical dataset (raw estimate: 0.563 g; standardised: 0.619, Supp Info S3). We acknowledge that we are assuming 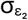 to be fixed even though it is likely that it also varies across contexts. Regardless, sensitivity analyses in which both residual values were varied across a plausible range confirmed that the slopes of the correlation reaction norm (β_r_), our primary inferential targets, were robust to these choices (table S1). However, variance reaction norm intercepts were conditional on the fixed residual values and should be interpreted accordingly.

Weakly regularising priors were used throughout to guard against overfitting while remaining permissive of biologically plausible effects, following Martin (2025): *Normal(0, 1)* for fixed-effect slopes and variance reaction norm slopes (β_σ_), *Normal(0, 0*.*5)* for correlation reaction norm slopes (β_r_), and *Exponential(2)* for all variance standard deviation parameters. The narrower prior on β_r_ reflects the expectation that very large shifts in the among-individual correlation across contexts are *a priori* unlikely, and to avoid placing undue weight on extreme correlations (Bliard et al., 2026). All models were implemented in a Bayesian framework using the statistical programming language Stan (Carpenter et al., 2017) via the R package *CmdStanR* (Gabry et al., 2025). Models in Part 1 and 2 each ran on three chains with a warm-up period of 1,000 iterations, sampling for a further 1,000 iterations per chain, yielding 3,000 posterior samples in total. The genetic CRN model in Part 3 (see below) was run with an extended warm-up period of 1,500 iterations followed by 3,000 sampling iterations per chain across three chains, yielding 9,000 posterior samples in total. Convergence was assessed by visual inspection of trace plots and by ensuring R-hat values were below 1.03 for all Part 1 and Part 2 parameters, and below 1.06 for all Part 3 parameters (Gelman & Rubin, 1992). Posterior predictive checks confirmed adequate model fit for both traits (Fig. S12). Throughout the results, we report posterior median effect sizes alongside 10–90% credible intervals.

### Part 1: Compensation CRN model

To characterise how the among-individual carry-over correlation varies simultaneously across natal and adult environments, we fitted the CRN model described above in which the context predictor matrix X_C_ included predictors from both life stages across three ecological axes: temperature, rainfall, and population density. We chose these three environmental axes as they are well-known drivers of body mass in great tits (López-Idiáquez, Cole, Satarkar, et al., 2026; Satarkar et al., 2026). Specifically, X_C_ contained an intercept, linear and quadratic terms for natal and adult temperature, and linear terms for natal and adult rainfall and natal and adult population density (9 predictors in total). Quadratic temperature terms were included because preliminary models restricted to temperature alone revealed credible nonlinearity in the correlation reaction norm. This is consistent with prior work in this population showing non-linear associations between temperature and body mass when temperature is measured over comparable individual-specific relative windows (López-Idiáquez, Cole, Satarkar, et al., 2026; Satarkar et al., 2026), which motivated testing whether a similar non-linearity might shape the carry-over correlation. Rainfall and density effects were adequately described by linear terms. Including all six environmental axes jointly means that each estimated slope reflects the independent contribution of that predictor to changes in the carry-over correlation.

### Part 2: Environmental mismatch CRN model

The CRN model in Part 1 estimates natal and adult environmental effects independently, but a complementary question is whether it is the degree of cross-life-stage mismatch (i.e., the divergence between the environment an individual developed in and the one it subsequently bred in), that drives the carry-over correlation rather than either environment in isolation. To test this, we re-parameterised X_C_ using mismatch scores computed as adult environment minus natal environment for each axis (temperature, rainfall, density), standardised prior to analysis. The context matrix included an intercept, and linear and quadratic mismatch terms for each of the three axes (7 predictors in total).

### Part 3: Genetic CRN model

To understand whether the phenotypic patterns identified in Parts 1 and 2 reflect genetic or non-genetic sources of individual variation, we extended the CRN to a bivariate animal model by incorporating the pedigree of the Wytham great tit population. The social pedigree was constructed from breeding records spanning 1960–2025 and comprised approximately 111,500 individuals. The Wytham great tit population has an extra-pair paternity rate of approximately 12% (Patrick et al., 2011), but simulations have shown that the social pedigree does not significantly bias quantitative genetic estimates, particularly for traits with low extra-pair paternity influence (Firth et al., 2015), and the Wytham social pedigree has been successfully used for heritability estimation of different traits in several recent studies (Jones et al., 2025; López-Idiáquez, Cole, Satarkar, et al., 2026; Satarkar et al., 2025). For model fitting, the pedigree was pruned to the 4,345 individuals analysed here and their ancestors using the *nadiv* package in R (Wolak, 2012), yielding a pruned pedigree of 8,633 individuals spanning 38 generations from which the numerator relatedness matrix was computed (the majority of pruned individuals are birds only recorded as nestlings and which left no descendants). All 4,345 focal individuals had both parents recorded in the pruned pedigree. The pedigree contained 2,734 parent-offspring pairs, 2,338 full sibling pairs, and 2,201 half-sibling pairs (1,250 maternal and 951 paternal), providing 7,273 informative relative pairs in total for estimating additive genetic variance.

The animal model uses the known relatedness structure among individuals to partition the total among-individual variance in each trait into an additive genetic component and a non-genetic individual component. In the CRN framework, both components had their own bivariate covariance matrix that was allowed to vary with environmental context. This yielded two separate correlation reaction norms estimated simultaneously; one describing how the genetic correlation between natal and adult body mass (r_A_) changes across environments, and one describing how the non-genetic individual correlation (r_E_) changes. This allowed us to directly ask whether the environmental modulation of the carry-over correlation identified in Parts 1 and 2 is driven by gene-by-environment interactions in the genetic covariance, or by environmentally varying non-genetic individual differences, or both.

We additionally computed narrow-sense heritability at mean environmental conditions, calculated for each posterior draw as h^2^ = V_A_ / (V_A_ + V_E_ + V_r_), using the reaction norm intercepts for additive genetic and non-genetic among-individual variance together with the fixed residual variance, before summarising the resulting posterior distribution as a median and 10%-90% credible intervals. However, because the nestling mass residual variance was fixed to reflect measurement error only, the h^2^ estimate should be interpreted cautiously. Although the shared within-brood environment among nestlings was accounted for through an explicit brood identity random effect, consistent maternal effects across broods were not separately modelled and may contribute to an upward bias in heritability estimates. Heritability at mean environment for adult mass is less affected by this issue, since its residual variance was independently estimated from repeated within-individual measures (see above). Heritability estimates are therefore best treated as approximate; our primary inferential targets were the reaction norm slopes of the genetic and non-genetic correlations, which are considerably less sensitive to the residual variance assumption than are absolute variance or heritability estimates.

The context matrix included an intercept, natal temperature, adult temperature, natal density, and adult density. Temperature and density were selected as the two focal axes because they were the most biologically motivated and empirically strongest modulators of the carry-over correlation in Parts 1 and 2. Linear terms were used throughout, as the primary question was whether r_A_ and r_E_ differ in their environmental sensitivity, which can be addressed with linear slopes; a supplementary model with quadratic temperature terms confirmed the main results were unchanged (Fig. S13 and S14). The genetic CRN slopes had lower effective sample sizes than their non-genetic counterparts, likely because r_A_ is estimated only through resemblance between relatives in the pedigree rather than the full individual-level dataset. To ensure sufficient posterior samples for these parameters, the model was run across three chains with an extended warm-up period of 1500 iterations followed by 3000 sampling iterations per chain, yielding 9000 total posterior draws.

## Results

### Part 1: Compensation CRN model

Under average environmental conditions, the among-individual correlation between nestling mass and adult body mass was r = 0.71 (posterior median, 10%–90% intervals = [0.665, 0.757]), indicating substantial developmental carryover across the population. All three environmental axes credibly modulated the strength of this correlation, though with varying degrees of confidence (Fig. 2; see Fig. 1 for guidance on interpreting reaction norm plots).

**Figure 2.**
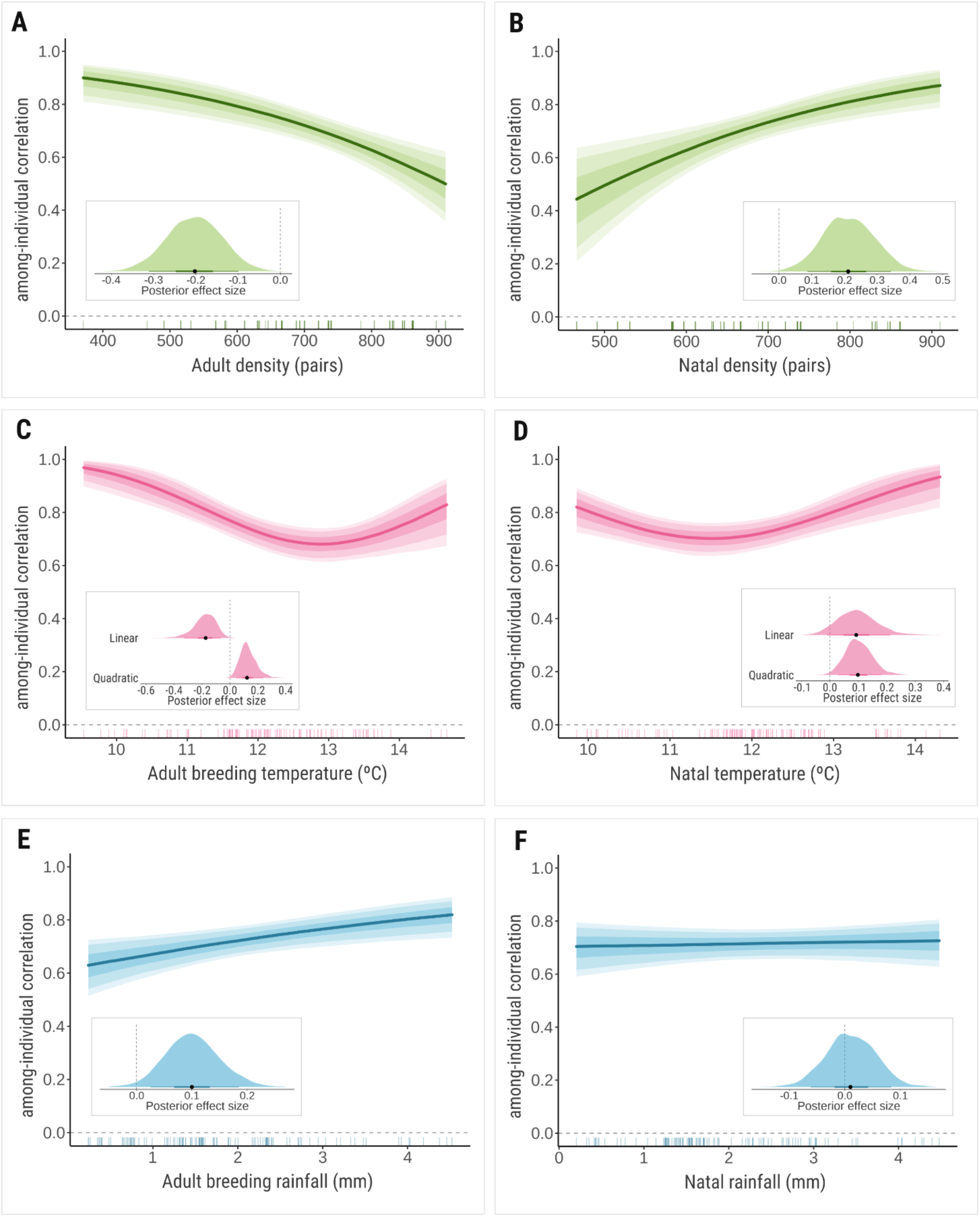
Among-individual correlations between nestling mass and adult body mass of great tits (*Parus major)* from Wytham Woods, UK as functions of (A) adult population density (number of pairs of great tits and blue tits *Cyanistes caeruleus* in the breeding year), (B) natal population density (number of pairs of great and blue tits in the hatch year), (C) adult temperature (average daily temperature in ºC during breeding), (D) natal temperature (average daily temperature in ºC during development i.e. 0–15 days post hatching), (E) adult rainfall (average daily rainfall in mm during breeding), (F) natal rainfall (average daily rainfall in mm during development i.e. 0–15 days post hatching); Lines show the posterior median effect sizes; shaded bands show 25–75%, 10–90%, and 5–95% credible intervals. Vertical lines at the x-axis show the distribution of observed context-level environmental values. Insets show the posterior distribution of the corresponding slope on the *atanh* scale (point = posterior median; thick bar = 50% CI; thin bar = 90% CI; dashed line = zero).

Population density produced the clearest effects on the natal-to-adult body mass correlation out of the three axes (Fig. 2A, 2B). Adult density had a robust negative effect (median = −0.202 [−0.286, −0.123]), with the correlation declining as the density of breeding great and blue tits increased (Fig. 2A). Natal density had an effect of similar magnitude but in the opposite direction (0.212 [0.115, 0.314]), with the correlation increasing across the observed density range (Fig. 2B).

Temperature showed non-linear effects at both stages (Fig. 2C, 2D). Adult breeding temperature had a robust quadratic effect (linear β = −0.174 [−0.285, −0.088]; quadratic β = 0.122 [0.059, 0.203]), with the credible intervals of both terms falling well clear of zero. The correlation decreased with increasing breeding temperatures, was weakest at warmer-than-average temperatures, and increased again at the extreme end (Fig. 2C). Natal temperature also showed a concave upward pattern (linear β = 0.093 [0.011, 0.183]; quadratic β = 0.098 [0.044, 0.161]), but the linear term’s lower bound was close to zero, and thus, is weakly supported, while the quadratic term is more clearly credible. The correlation was weakest at slightly cooler-than-average natal temperatures, and strengthened more steeply towards warmer extremes (Fig. 2D). However, given the marginal linear term, this shape should be interpreted cautiously, particularly regarding the weakest point of the correlation.

Adult rainfall had a credible positive effect (median = 0.100 [0.041, 0.164]), with the correlation rising across the observed rainfall range (Fig. 2E), albeit with a smaller effect size than density or temperature. However, natal rainfall showed no detectable effect (0.010 [−0.047, 0.067]) on the among-individual correlation between natal and adult body mass, which remained stable across the observed range (Fig. 2F).

Variance reaction norms were estimated jointly with the correlation reaction norms for both traits across all environmental axes; full results are reported in Supplementary Info S4.1. Fixed effects on trait means all followed expected directions (Fig. S10). Lay date, natal brood size, natal rainfall, and phenological mismatch, all credibly reduced chick weight; breeding temperature and number of offspring credibly reduced adult mass, while age and sex (male) predicted higher adult mass. Both birth year and breeding year showed credible negative linear trends (birth year β = −0.235 [−0.290, −0.182]; breeding year β = −0.214 [−0.250, −0.177]), indicating that both nestling and adult body mass have declined over the study period, consistent with a recent study of long-term body mass trends in this population (López-Idiáquez, Cole, Satarkar, et al., 2026).

### Part 2: Environmental mismatch CRN model

The among-individual correlation between nestling and adult body mass as a function of the mismatch between natal and adult temperatures was a U-shaped curve, with both linear and quadratic terms for thermal mismatch credible (linear β = −0.205 [−0.329, −0.110]; quadratic β = +0.187 [0.106, 0.294]). The correlation was weakest when natal and adult thermal environments were similar, and strengthened as the two diverged in either direction (Fig. 3B). The minimum of the curve, however, was not at zero mismatch (i.e., natal temperature = adult temperature), but when adult temperatures were slightly warmer than natal temperatures.

**Figure 3.**
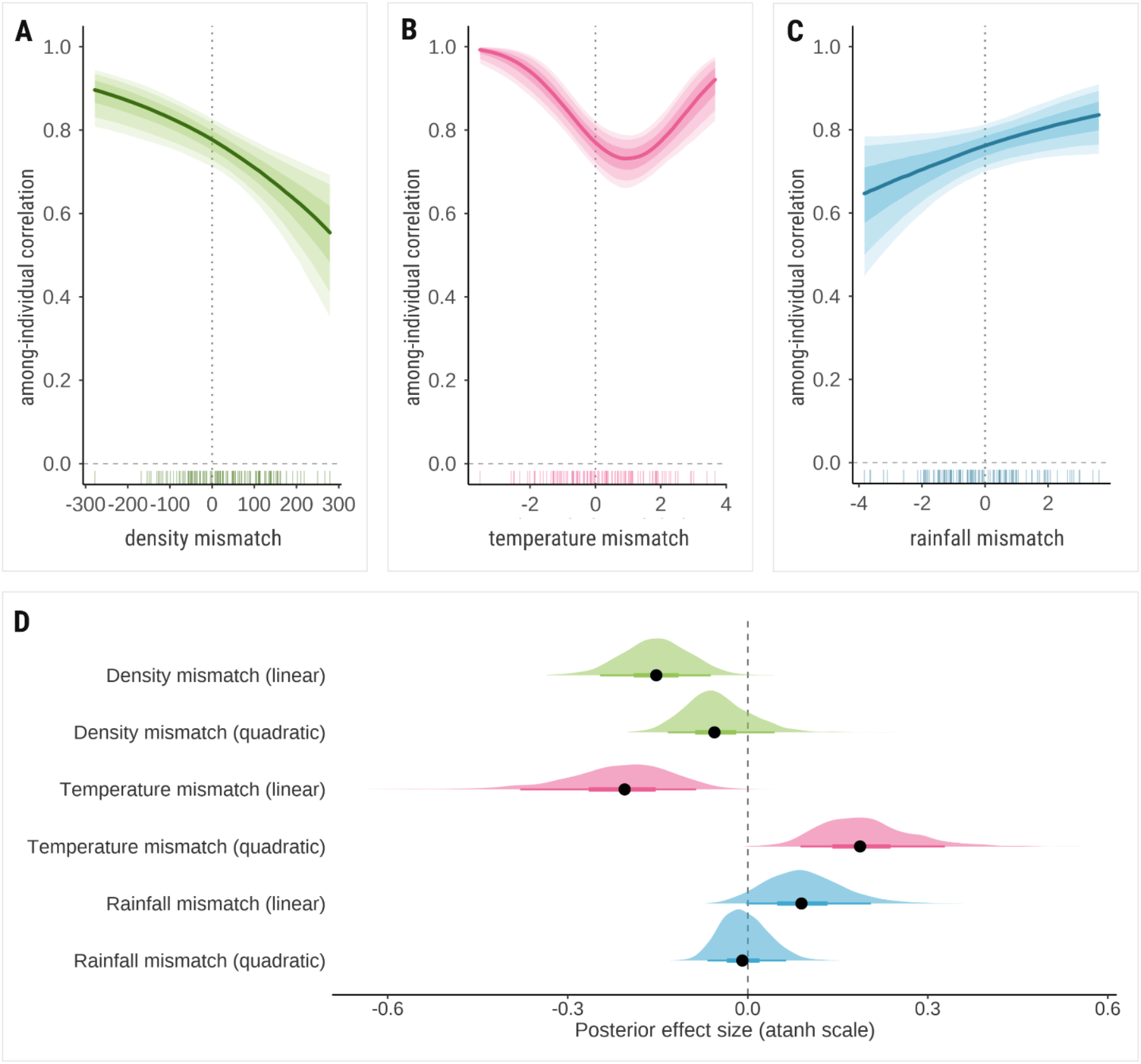
Among-individual correlations between nestling mass and adult body mass of great tits (*Parus major)* from Wytham Woods, UK as functions of mismatch (calculated as difference) between; **(A)** adult population density (number of pairs of great tits and blue tits *Cyanistes caeruleus* in the breeding year) and natal population density (number of pairs of great and blue tits in the hatch year); **(B)** adult temperature (ºC) (average daily temperature during breeding), and natal temperature (ºC) (average daily temperature during development (0–15 days post hatching)); **(C)** adult rainfall (mm) (average daily rainfall during breeding), and natal rainfall (mm) (average daily rainfall during development (0–15 days post hatching)). Lines show the posterior median effect sizes; shaded bands show 25–75%, 10–90%, and 5–95% credible intervals. Vertical lines at the x-axis show the distribution of observed context-level environmental values. **(D)** Posterior distributions of the corresponding CRN slopes on the *atanh* scale (point = posterior median; thick bar = 50% CI; thin bar = 90% CI; dashed line = zero).

Mismatch in natal and adult population density showed a credible negative effect (median = −0.152 [−0.225, −0.081]). The correlation was high when natal density exceeded adult density, but weakened as adult density increasingly exceeded natal density (Fig. 3A), consistent in direction with the density effects identified in Part 1. Mismatch in natal and adult rainfall showed a weakly credible positive linear effect (0.089 [0.015, 0.177]); the correlation strengthened as adult rainfall increasingly exceeded natal rainfall (Fig. 3C).

Fixed effects on trait means closely matched those from Part 1 (Fig. S10), as expected given the same predictors for both traits. Variance reaction norms were jointly estimated with the correlation reaction norms as before (Supp. Info S4.2).

### Part 3: Genetic CRN model

We were able to partition the total phenotypic carry-over correlation into a high additive genetic correlation (r_A_ = 0.861 [0.745, 0.931]) and a substantial non-genetic among-individual correlation (r_E_ = 0.768 [0.661, 0.859]) at mean environmental conditions. Heritability at this baseline was h2 = 0.474 [0.373, 0.568] for nestling mass and h^2^ = 0.349 [0.278, 0.411] for adult body mass. These estimates are broadly consistent with previous heritability estimates of body mass in this population, despite differing modelling approaches (Garant et al., 2004; López-Idiáquez, Cole, Satarkar, et al., 2026).

None of the four environmental slopes for r_A_ were credible (natal temperature β = 0.098 [−0.160, 0.330]; adult temperature β = 0.164 [−0.102, 0.424]; natal density β = −0.179 [−0.554, 0.212]; adult density β = −0.011 [−0.445, 0.343]) (Fig. 4). Given the width of the intervals, all overlapping zero, this indicated no evidence of gene-by-environment interactions for the natal-to-adult body mass link over the observed range of environments in this population.

**Figure 4.**
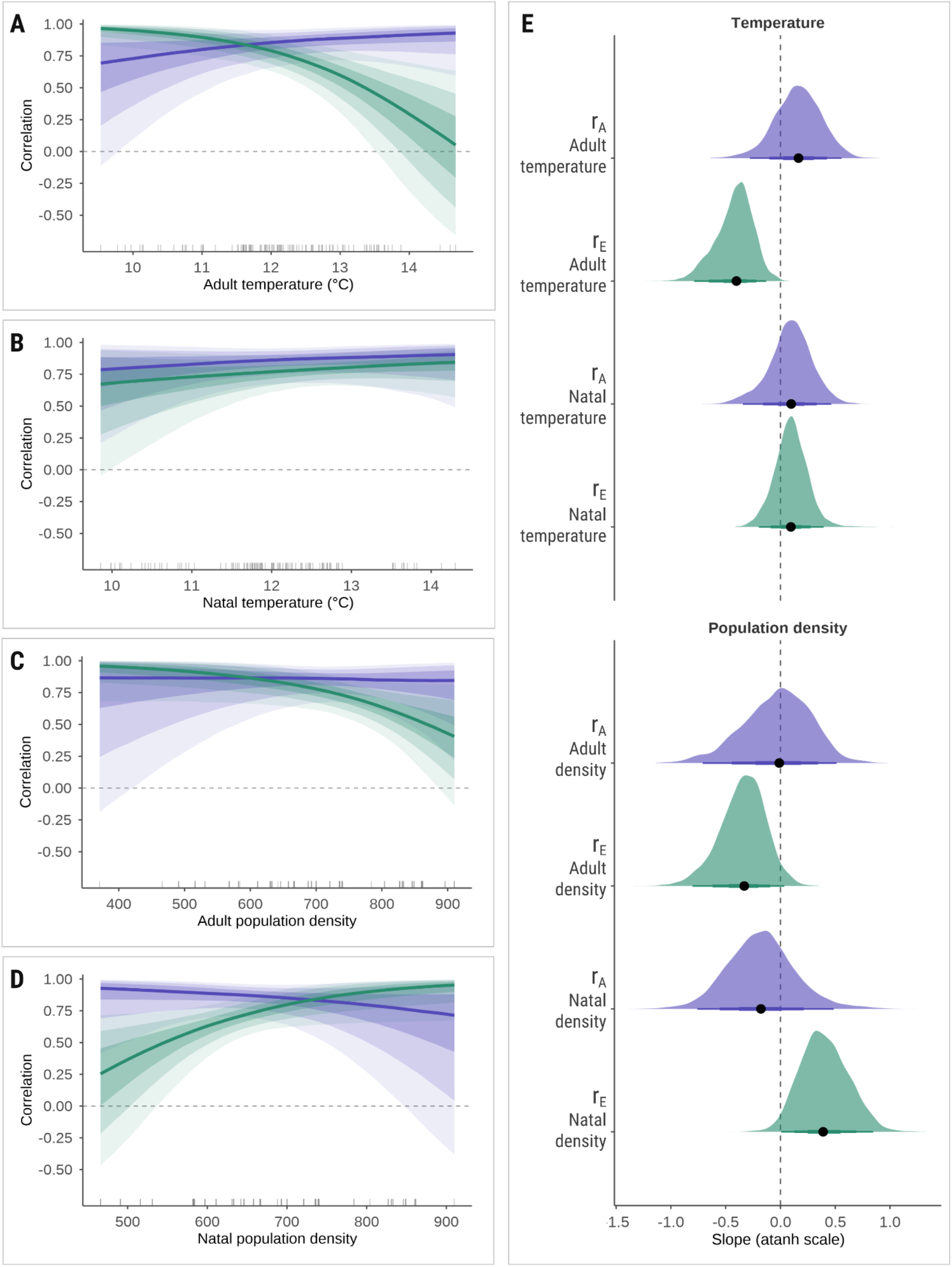
Reaction norms for the additive genetic correlation (r_A_ in purple) and non-genetic among-individual correlation (r_E_ in green) between nestling mass and adult body mass of great tits (*Parus major)* from Wytham Woods, UK as functions of **(A)** adult temperature (ºC) (average daily temperature during breeding), **(B)** natal temperature (ºC) (average daily temperature during development (0–15 days post hatching)), **(C)** adult population density (number of pairs of great tits and blue tits *Cyanistes caeruleus* in the breeding year), **(D)** natal population density (number of pairs of great and blue tits in the hatch year), Lines show the posterior median effect sizes; shaded bands show 25–75%, 10–90%, and 5–95% credible intervals. Vertical lines at the x-axis show the distribution of observed context-level environmental values. **(E)** Posterior distributions of the corresponding CRN slopes on the *atanh* scale (point = posterior median; thick bar = 50% CI; thin bar = 90% CI; dashed line = zero).

In contrast, r_E_ was credibly modulated by adult breeding temperature, and population density during both natal and adult life stages (adult temperature β = −0.402 [−0.655, −0.220]; natal density β = +0.389 [0.127, 0.692]; adult density β = −0.331 [−0.620, −0.094]). The patterns for r_E_ directly mirror the environmental modulation of the phenotypic correlation identified in Part 1 on the same axes, indicating that the among-individual phenotypic correlation between nestling and adult body mass can be largely attributed to non-genetic individual differences, rather than a genetic basis.

Although r_A_ itself showed no credible environmental modulation, the genetic variance underlying nestling mass did. All four environmental slopes for the nestling weight genetic variance reaction norm were credible (natal temp β = −0.283 [−0.442, −0.068]; adult temp β = +0.218 [0.055, 0.374]; natal density β = +0.210 [0.014, 0.399]; adult density β = −0.211 [−0.389, −0.013]; Fig. S9). For adult body mass, only the adult temperature slope on genetic variance was marginally credible (β = +0.246 [0.025, 0.434]); the natal density and adult density slopes showed poor chain mixing and were excluded from interpretation (Fig. S9).

## Discussion

Using a covariance reaction norm model (Martin, 2025) and 40 years of individual-based data from one of the longest-monitored populations of wild birds (Perrins, 1965), we show that developmental carry-over of body mass in the Wytham great tit population is strong, but not fixed. We show that the association between natal and adult body mass varies continuously with the environmental conditions experienced at both life stages, and that this sensitivity results almost entirely from the non-genetic component of the correlation rather than the genetic one. Our results, therefore, reframe the ‘silver-spoon’ effect, not as a fixed developmental outcome, but as a context-dependent relationship whose strength depends on the ecological conditions an individual experiences across its life.

Population density was the strongest modulator of developmental carry-over strength, with the correlation declining markedly as adult density increased and strengthening when natal density was higher. This pattern fits well with classic ideas about density-dependent processes in great tits. In Wytham specifically, local breeding density has been shown to influence fledging mass and recruitment (Wilkin et al., 2006), with lighter fledglings produced when females bred in smaller territories (i.e., at higher density). Heavier fledglings have also been shown to gain a particular advantage in recruitment under high density conditions (Both et al., 1999). More recently, it has been shown in the Wytham population that increasing density is linked to declines in both nestling and adult mass, with effects carrying across life stages (López-Idiáquez, Cole, Satarkar, et al., 2026). Our finding that density modulates the natal-to-adult body mass relationship itself, rather than just the mean level of each trait, can help shed light on plausible mechanisms underlying the carry-over of body mass across life stages.

When breeding density is high, resources are likely constrained for all birds, so even individuals that began life in better condition may be unable to translate that early advantage into higher adult mass, weakening the correlation. By contrast, when adult breeding density is low, resources are abundant, allowing individuals with an early-life advantage, for instance having been heavier as nestlings, to capitalise on favourable conditions and maintain a stronger natal-to-adult association. Alternatively, high breeding density may itself be a downstream signal of favourable conditions in the preceding year. Breeding density in this population is strongly influenced by abundance of beech mast and overwinter survival (Perrins, 1965). In this scenario, benign, favourable conditions would allow even poor-quality individuals to attain typical adult mass through compensatory growth, weakening the correlation regardless of direct resource competition among adults. Since we are interpreting correlations, it is challenging to distinguish between these explanations. However, nestling mass is considerably more sensitive to breeding density than adult mass in this population (López-Idiáquez, Cole, Satarkar, et al., 2026), suggesting that nestlings may be bearing a disproportionate cost of high density as opposed to adults. At the natal density level, competitive natal environments appear to sharpen the signal of individual quality, so that only the highest-quality nestlings maintain body mass under resource limitation, making natal mass a more informative predictor of adult mass, and implying a stronger correlation. This interpretation is also supported by the increase in nestling mass variance under high natal density (Fig. S5), which suggests that competitive developmental conditions amplify differences among individuals rather than simply reducing growth uniformly across the cohort.

Temperature and rainfall effects on the body mass correlation may also be operating through compensatory mechanisms. Breeding temperature showed a nonlinear relationship with the correlation, which was weakest at intermediate-to-warm temperatures and strongest at the extremes. The initial decline in carry-over strength fits a compensatory explanation well (Metcalfe & Monaghan, 2001): benign, warm, conditions may give individuals that started life in poorer condition an opportunity to catch up, narrowing the gap with those that had an early-life advantage and therefore weakening the link between nestling and adult mass. By contrast, wetter breeding conditions, which are more constraining for foraging and provisioning, strengthen the correlation, consistent with harsher adult conditions limiting the capacity of lower-quality individuals to compensate and allowing early-life differences to persist. The increase in carry-over at the warmest temperatures may reflect a related but distinct process, whereby extreme heat itself becomes a stressor that only higher-quality individuals can tolerate and use to maximise their natal carried-over quality. Natal temperature and rainfall, by contrast, had only weak or no detectable effects on the correlation, suggesting that it is primarily the weather conditions individuals experience as adults, rather than those they experienced as nestlings, that determine how much of an early-life advantage is allowed to persist.

This pattern held true even when the resemblance between natal and adult environments was considered. Thermal mismatch produced a U-shaped curve, with carry-over strength weakest when natal and adult temperatures were similar, and strengthening as they diverged in either direction. This is contrary to our initial expectation, as pure environmental matching assumes that any matched pair of environments should produce a strong correlation, simply because they match, regardless of whether that happens to be cold-cold or warm-warm. However, our results suggest that ‘matched’ environments need not produce the same correlation because the adult environment itself strongly drives how much of the natal advantage is retained. The environmental matching hypothesis is usually applied to fitness consequences or individual performance, but our results highlight its limitations when applied to a correlation. In Part 2, when adult conditions were harsher than natal ones (e.g., breeding temperatures colder than those experienced as a nestling), early-life differences persisted and the correlation remained strong; when adult conditions were more benign than natal ones (e.g., slightly warmer breeding conditions), individuals could compensate and the correlation weakened. Rainfall and density mismatch showed weaker, more directional patterns that largely mirrored their respective adult-stage effects from Part 1, reinforcing that it is the adult environment, not the degree of resemblance to the natal one, that carries the most weight. This is particularly relevant under continued climate change, which is projected to increase not just mean temperatures but increased variability in temperature and rainfall (IPCC, 2023). As natal and adult environments become more likely to diverge within a single lifetime, our results suggest that carry-over may not depend simply on whether natal and adult environments match, but on the direction of that mismatch and on how restrictive or benign adult conditions are.

We showed that the genetic correlation (rA) between nestling and adult mass was stable across the thermal and density gradients we tested, while the non-genetic component of the among-individual correlation (rE) shifted with the environment in the same directions as the phenotypic patterns described above. This correlation-level finding is analogous to what has been established for labile traits, which is that phenotypic plasticity can vary systematically with the environment via non-genetic sources without underlying gene-by-environment interactions (Nussey et al., 2007). Furthermore, while genetic correlations are not fixed properties of trait pairs, and may shift, or even reverse, across environments, they can be more resistant to such changes when traits are tightly bound by shared physiology or development (Stearns et al., 1991). Nestling and adult body mass are likely expressions of a tightly-linked genetic architecture reflecting an individual’s growth trajectory, and thus, we can reasonably expect an environmentally-stable genetic correlation among these two traits. However, a stable genetic correlation does not necessarily mean the genetic consequences of selection are themselves stable. The genetic variance underlying nestling mass varied credibly across all four environmental axes we tested, despite the correlation being fixed. This suggests that environmental conditions at different life stages are shaping genetic variance within each trait without changing how tightly the traits are genetically linked, while strongly modulating the non-genetic variance-covariance structure. Therefore, our results indicate that climate change and population density are altering how strongly early-life advantages are expressed, buffered, or lost in the phenotype, and how much scope there is for selection to act on them, without necessarily reconfiguring the genetic architecture underpinning body mass across life stages.

Our study has an inherent caveat in its design, in that developmental carry-over is measured only among individuals that survive to adulthood and thus, estimated variances and correlations are conditional on prior viability selection. The ‘missing fraction’ problem describes how selection on a trait correlated with a later-life trait can distort the covariance between them, because the individuals measured are a non-random subset of the original cohort, filtered by selection (Mittell & Morrissey, 2024). Therefore, the environment-dependent patterns we report in the natal-to-adult body mass correlation are likely shaped, in part, by selection on the nestlings that went on to breed, rather than purely by processes acting after nestlings recruit. This is particularly plausible given that selection on nestling mass in this population has increased over the study period, and can also be density-dependent (Both et al., 1999; López-Idiáquez, Cole, Satarkar, et al., 2026). Nevertheless, this limitation does not critically undermine our main conclusions, as our aim was to characterise how environment shapes carry-over among the individuals that actually go on to contribute to the next generation. An important future direction would be to model survival as a third trait, such that genetic covariance can be estimated via resemblance among relatives, allowing us to determine how much of the observed environmental dependence arises before recruitment versus after it, without requiring adult-mass phenotypes for non-surviving individuals. As climate change induces rapid environmental variation, disentangling whether developmental advantages are being reshaped by changing environments or changing selection regimes will be important for determining how reliably early-life body mass can be used to infer individual quality, and for predicting its evolutionary trajectory in wild populations.

## Acknowledgements

We thank the numerous researchers and fieldworkers involved in the collection of the long-term data for the Wytham tit study over the past 80years. The long-term population study has been supported by numerous funding sources, including recently by grants from BBSRC (BB/L006081/1), NERC (NE/K006274/1, NE/S010335/1) and a UKRI Frontiers Award (EP/X024520/1), together with NERC award NE/X000184/1 which partially supported this work

## Author contributions

Devi Satarkar conducted the statistical analyses with substantial inputs on methodology from Louis Bliard and David López-Idiáquez. All authors contributed to the ideas of this study. Devi Satarkar drafted the manuscript with all authors providing critical feedback. All authors approved the final manuscript.

## Data and code availability

All the data, R and Stan code needed to reproduce the analyses and figures in this manuscript are available at https://github.com/devisatarkar/CRN_bodymass_Wytham

