## Supplementary Information for "Strength of developmental carry-over varies continuously with environmental conditions in a wild bird population"

#### S1. Defining contexts

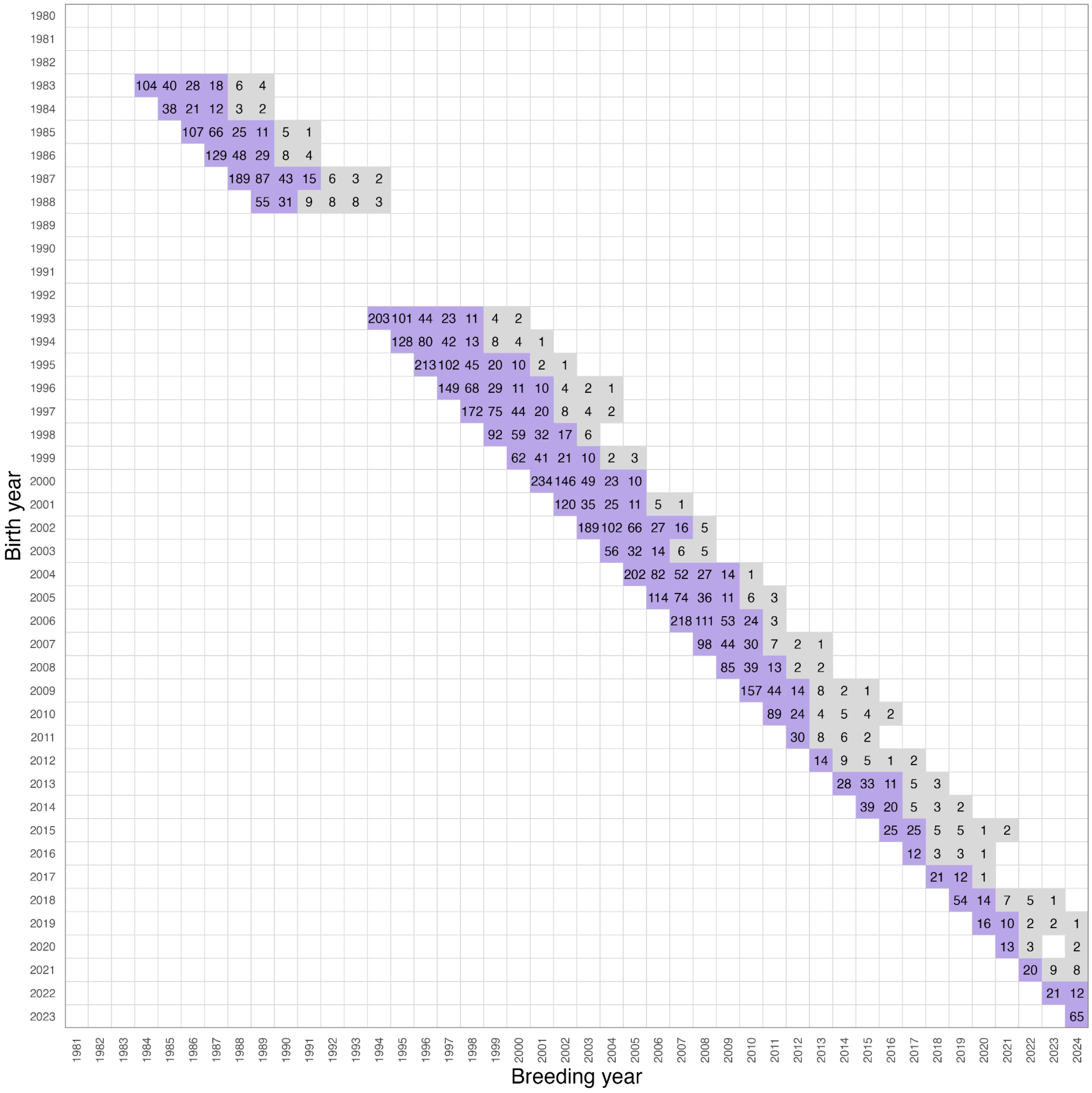


**Figure S1.** Sample sizes for each birth year x breeding year context. Each cell represents the number of individuals that hatched in the corresponding birth year, and went on to breed in the corresponding breeding year. Purple cells are the contexts retained for analysis (113 contexts, N ≥ 10 individuals),

Grey cells are excluded contexts (N<10); blank cells indicate no individuals present in the context for our dataset. Please note that this context matrix corresponds to the final dataset used for analysis, after filtering for complete cases across all variables.

Environmental variables were context-specific, and corresponded to the overall natal and adult environments experienced by individuals belonging to a context. For example, the 65 individuals that hatched in 2023, and bred in 2024 (bottom-right corner cell), each had individual-level natal temperature, adult temperature, natal rainfall, and adult rainfall values, corresponding to their developmental periods (0-15 days old), and the developmental periods of their own offspring in their breeding attempt (0-15 days old offspring). The context-level variables for the 2023 x 2024 context were calculated as the mean of the 65 natal temperature, adult temperature, natal rainfall, and adult rainfall values. Therefore, context-level climate variables were means of means, reflecting the actual thermal and precipitation conditions experienced by the specific cohort of individuals in that context, rather than a generic seasonal average for that year. Population density was assigned to each year directly, corresponding to the total number of breeding pairs of great tits and blue tits in a given year.

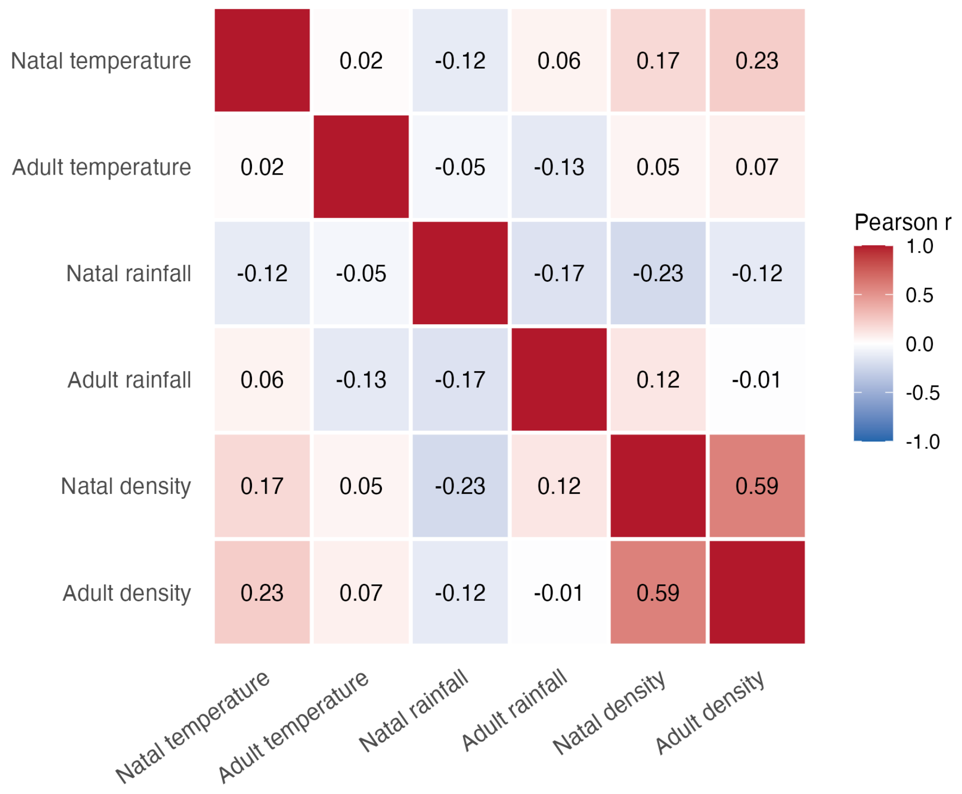


**Figure S2.** Correlations among the predictor variables used in the model, with low to moderate collinearity among all variables.

### S2. Missing data imputation (Sensitivity analysis for reduced samples)

As Fig. S1 shows, we did not have records for individuals between 1989–1992. These individuals were dropped while filtering for complete cases, as we do not have the annual winter moth (*Operophtera brumata*) half-fall date estimates for these years. The natal phenological mismatch variable, which is used as a fixed effect for nestling mass in all the three models, is calculated as the absolute difference between the half-fall date of winter moth larvae and the tenth day post-hatching of the focal individual.

As a robustness check, we imputed the unknown half-fall date values using predictive mean matching with the R package *mice* (ref), and ran the Part 1 models on 5 versions of the imputed datasets derived from these recovered missing values, in order to account for the uncertainty associated with imputation. Recovering these years increased our sample sizes from 6,686 to 8,717 breeding attempts by 5,851 individuals (previously 4,447), and the number of environmental contexts with N<10 from 113 to 149. Posterior estimates from the analysed imputed datasets were pooled and compared against the primary analysis as reported in the main text (Part 1).

CRNs for density and adult temperature were directionally unchanged and remained credible, although adult density and the adult temperature quadratic term were slightly attenuated in magnitude. Adult rainfall remained credibly positive, while natal rain was not credible in both analyses. The effects of the natal temperature terms (linear and quadratic) remained marginal as in the primary analysis, but revered sign in the imputed version, with the quadratic term no longer credibly differing from zero. However, our primary analysis already treated the natal temp CRN slope cautiously, given the marginal linear term (Fig. 2D). All other reported effects and the overall conclusions of the analyses reported in the main text are largely unaffected by this sensitivity check using imputed data, and our reduced sample sizes (due to 4 years of missing data; Fig S1) are sufficient to detect environmental modulation of correlations between nestling and adult body mass (see Fig 2 and Fig S3 for comparison).

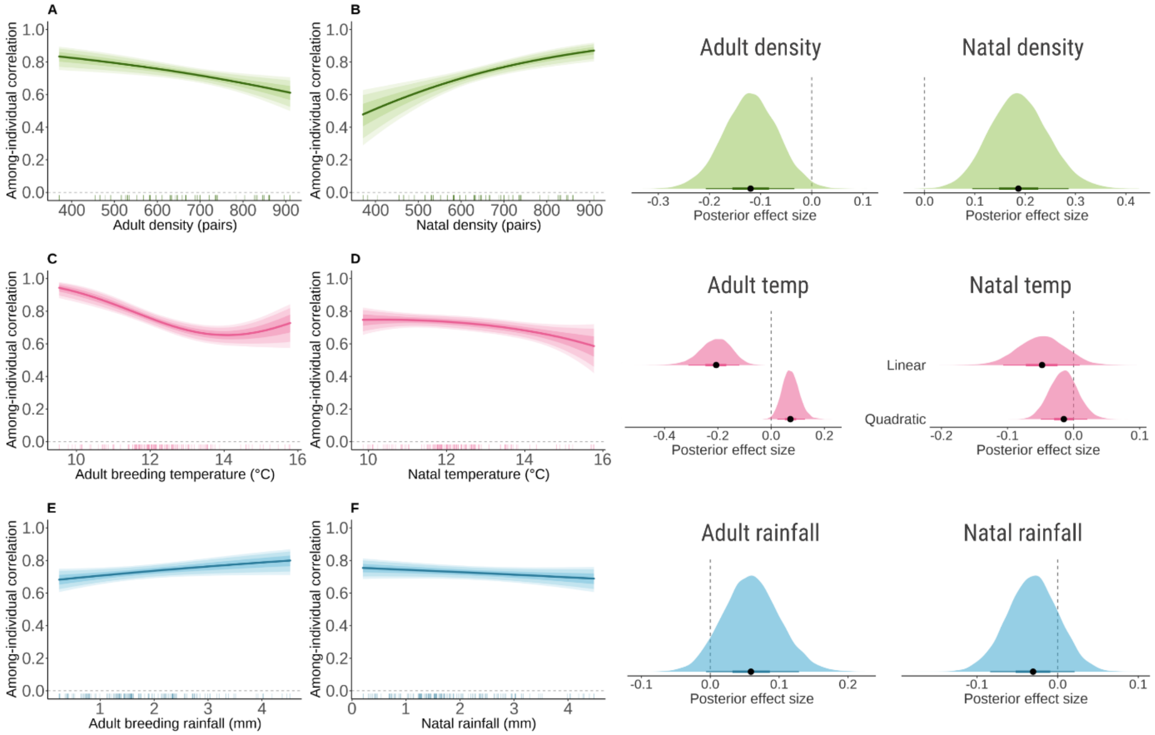


**Figure S3.** Among-individual correlations between nestling mass and adult body mass of great tits from Wytham Woods, UK as functions of **(A)** adult population density, **(B)** natal population density, **(C)** adult temperature, **(D)** natal temperature, **(E)** adult rainfall, **(F)** natal rainfall using imputed missing data; Lines show the posterior median effect sizes; shaded bands show 25–75%, 10–90%, and 5–95% credible intervals. Vertical lines at the x-axis show the distribution of observed context-level environmental values. Panels on the right show the posterior distribution of the corresponding slope on the *atanh* scale (point = posterior median; thick bar = 50% CI; thin bar = 90% CI; dashed line = zero). Posterior densities are for 15,000 pooled draws (3000 draws per model; 5 models corresponding to 5 versions of imputed datasets for missing half-fall dates).

### S3. Fixed residual standard deviations

The residual standard deviation (SD) for adult mass ($\sigma_{\varepsilon_2}$) was derived from a univariate linear mixed model fitted to the same analytical dataset using the R package *lme4*. The response variable was adult mass, and the fixed effects were the same as used for the primary CRN models reported in the main text (laying date, mean temperature, mean rainfall, number of offspring, age, sex, and breeding year). The random effects were individual identity, nestbox and breeding year. This yielded a repeatability of 0.502 for adult mass, and a residual SD of 0.619 on the standardised scale, which was fixed as $\sigma_{\varepsilon_2}$ for the CRN models (Part 1-3). The residual SD for nestling mass ($\sigma_{\varepsilon_1}$) was fixed as 0.10, a value broadly consistent with the known precision of the scales used to weigh nestlings. reflecting the standard measurement precision of weighing scales. Since nestling mass is measured only once, we cannot directly estimate within-individual variance for this trait, and the residual and among-individual variance cannot be distinguished from the data alone. In these situations, the residual term may be fixed based on external knowledge (Ponzi et al., 2018). The magnitude of $\sigma_{\varepsilon_1}$ can directly affect among-individual variance and heritability estimates. Smaller values shift more of the total phenotypic variance into the among-individual component, in turn inflating heritability.

We tested the sensitivity of the CRN and VRN slopes to plausible alternative values, with $\sigma_{\varepsilon_1}$ = 0.05 and 0.2, and $\sigma_{\varepsilon_2}$ = 0.3 and 0.9 (Table S1). This check was conducted using an earlier CRN model specification restricted to the linear term for temperature only. The CRN slopes for natal and adult temperature were robust across the range of values used for the residual SDs, except $\sigma_{\varepsilon_2}$ = 0.9. However, we did not find this alarming, because 0.9 is an implausibly high value for within-individual variation in adult mass.

**Table S1.** Sensitivity of the temperature CRN and VRN slopes to fixed residual standard deviations for nestling mass and adult mass. Values are posterior median slope estimates (CRN slopes on *atanh* scale, and VRN on *log* scale). P(neg) or P(pos) gives the posterior probability that the slope is in the stated direction. Bold rows indicate the residual values used in the primary analyses reported in the main text.

| **Nestling mass residual SD $\sigma_{\varepsilon_1}$** | **CRN – adult temp** | **P(neg)** | **CRN – natal temp** | **P(pos)** | **VRN – adult temp** | **P(pos)** |
|---|---|---|---|---|---|---|
| 0.050 | −0.156 | 1.000 | +0.067 | 0.952 | +0.136 | 1.000 |
| **0.100** | **−0.162** | **1.000** | **+0.067** | **0.943** | **+0.136** | **1.000** |

| | | | | | | |
|---|---|---|---|---|---|---|
| 0.200 | −0.171 | 1.000 | +0.063 | 0.930 | +0.136 | 1.000 |
| **Adult mass residual SD $\sigma_{\varepsilon_2}$** | **CRN – adult temp** | **P(neg)** | **CRN – natal temp** | **P(pos)** | **VRN – adult temp** | **P(pos)** |
| 0.300 | −0.069 | 1.000 | +0.035 | 0.948 | +0.086 | 1.000 |
| **0.619** | **−0.162** | **1.000** | **+0.067** | **0.943** | **+0.136** | **1.000** |
| 0.900 | −0.103 | 0.686 | −0.011 | 0.486 | +0.073 | 0.862 |

### S4. Variance reaction norms for nestling mass and adult body mass

#### S4.1. Part 1: Compensation CRN model

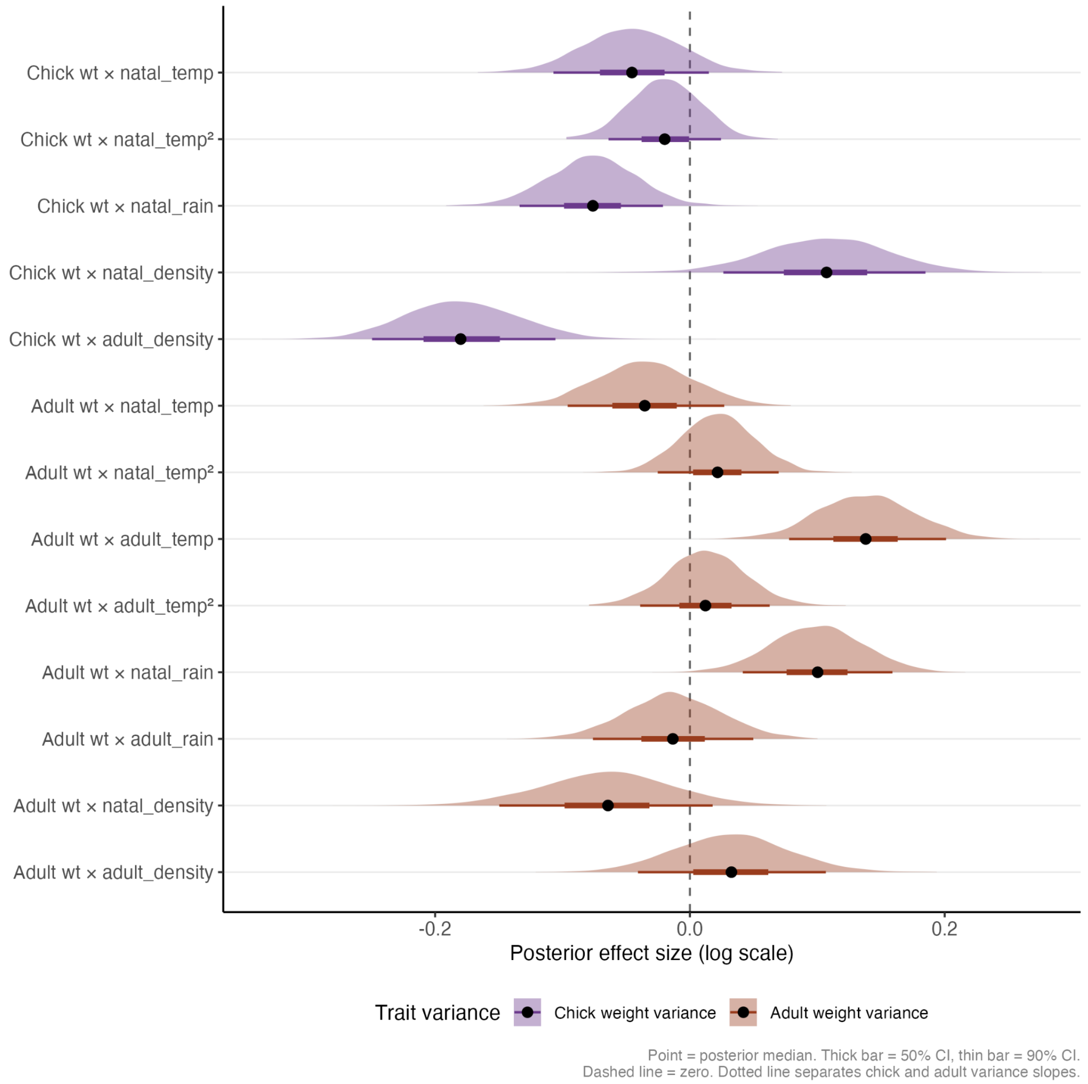


**Figure S4.** Posterior densities for variance reaction norms for nestling mass (labelled Chick wt) and adult mass (Adult wt) across temperature, rainfall and density axes

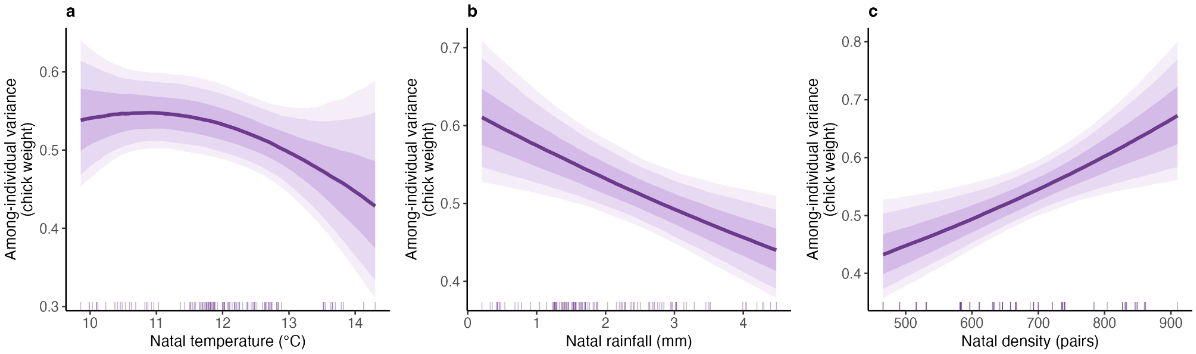


**Figure S5.** Variance reaction norms for nestling mass along natal environmental gradients (only rain and density credible).

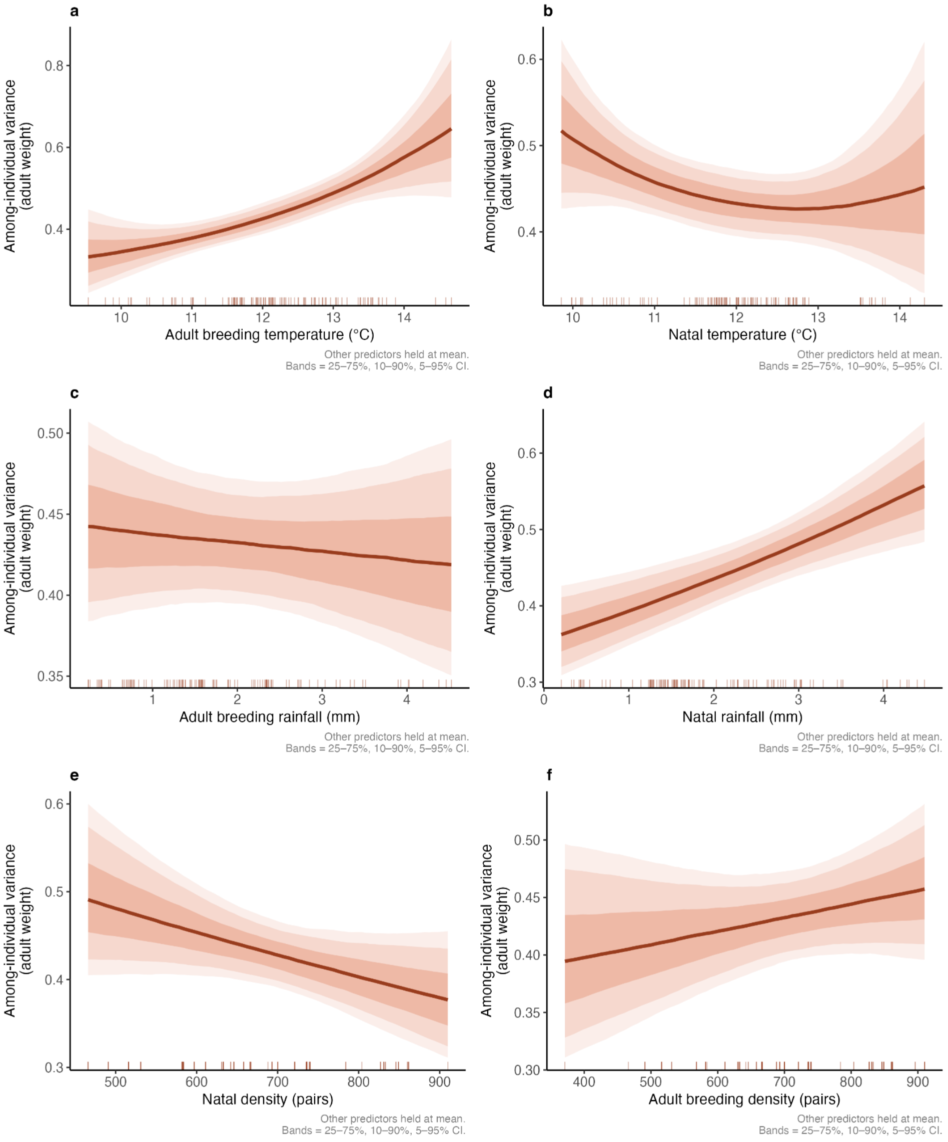


**Figure S6.** Variance reaction norms for adult mass along natal and adult environmental gradients

### S4.2. Part 2: Environmental mismatch CRN model

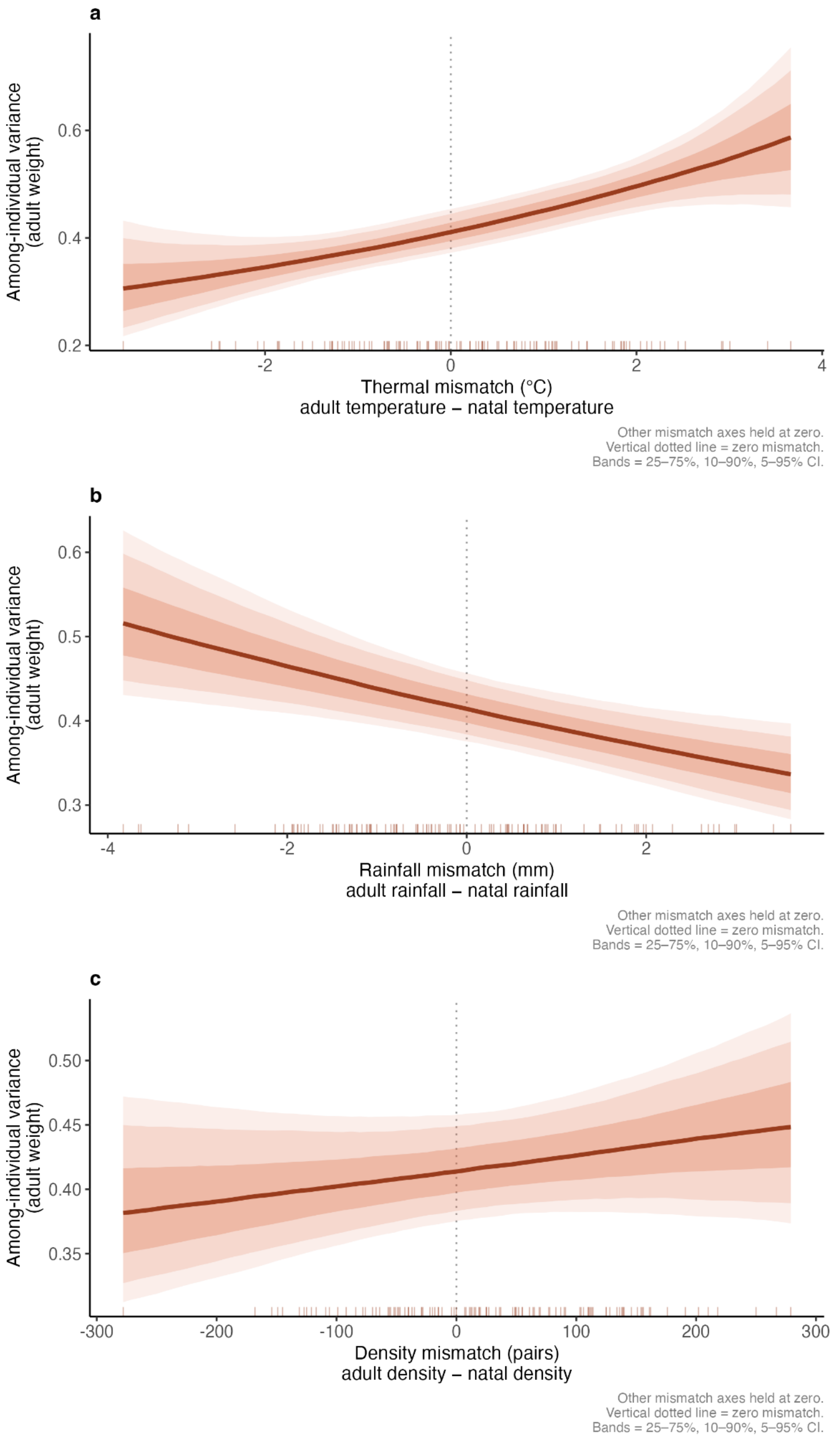


**Figure S7.** Variance reaction norms for adult mass along environmental mismatch gradients i.e. adult environment - natal environment.

#### S4.3. Part 3: Genetic CRN model

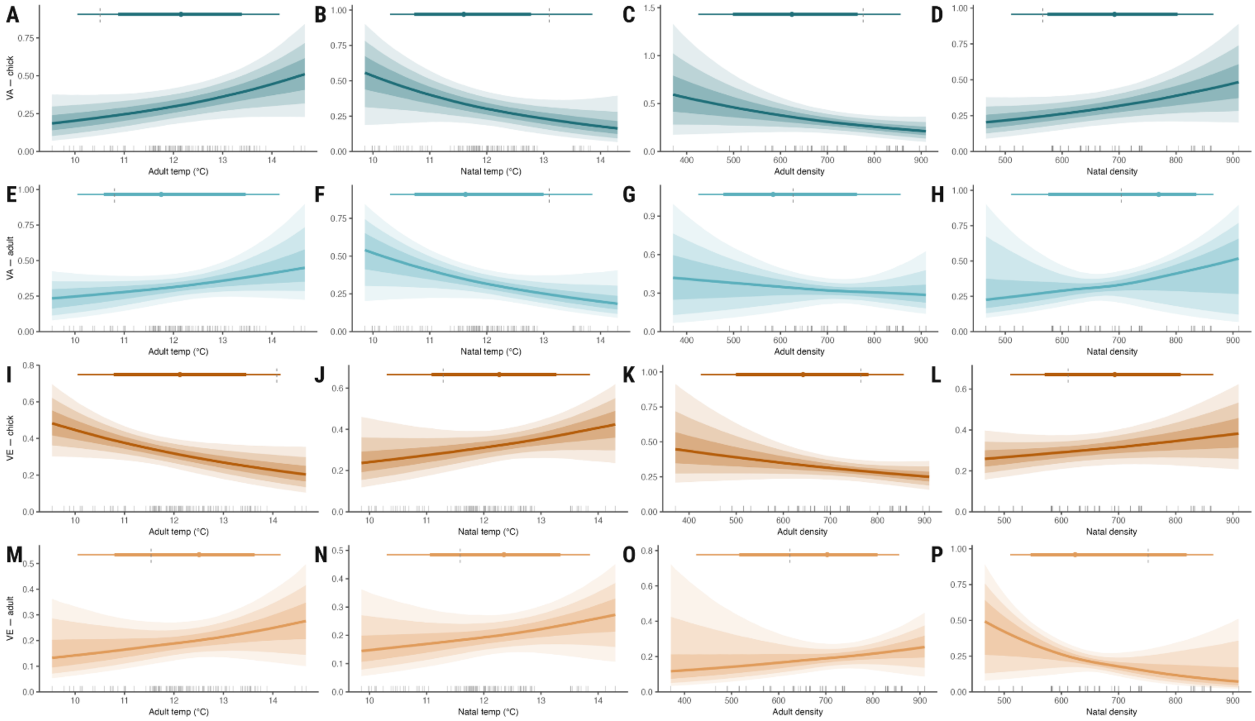


**Figure S8.** Variance reaction norms of genetic (A-H) (blue) and non-genetic (I-P) (orange) components for nestling mass (A-D, I-L) (VA – chick, VE – chick) and adult mass (E-H, M-P) (VA – adult; VE – adult) along environmental gradients: Adult temperature (in ºC), natal temperature (in ºC), adult population density, and natal population density. Insets show the posterior medians, with thick bar = 80 % credible intervals, and thin bar = 95% credible intervals, relative to zero (grey dotted line).

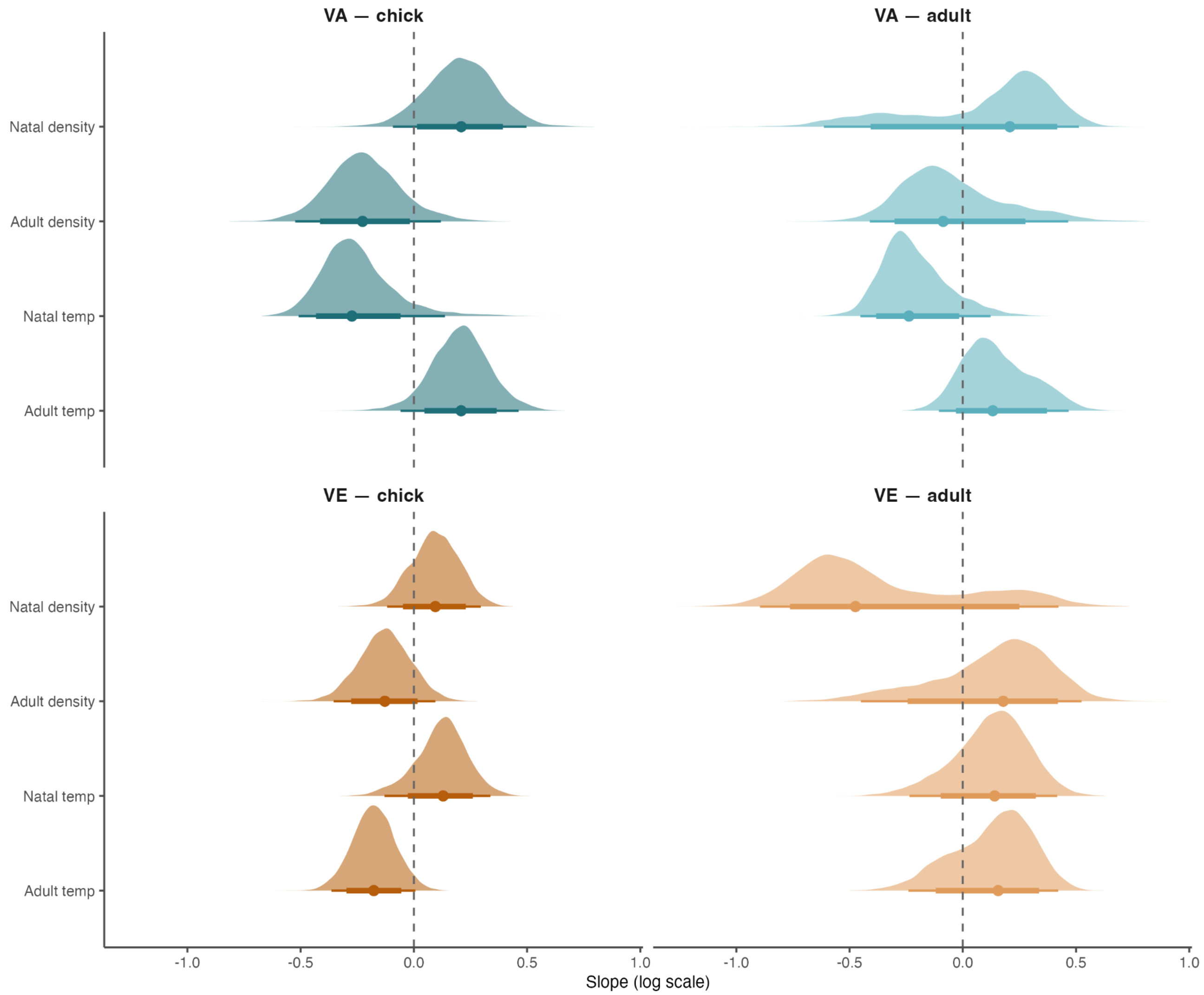


**Figure S9.** Posterior densities for variance reaction norms of genetic (VA; blue) and non-genetic (VE; orange) components for nestling mass (VA – chick, VE – chick) and adult mass (VA – adult, VE – adult) across natal and adult temperature and density axes.

### S5. Fixed effects on nestling mass and adult mass

#### S5.1 Part 1 and Part 2 CRN models

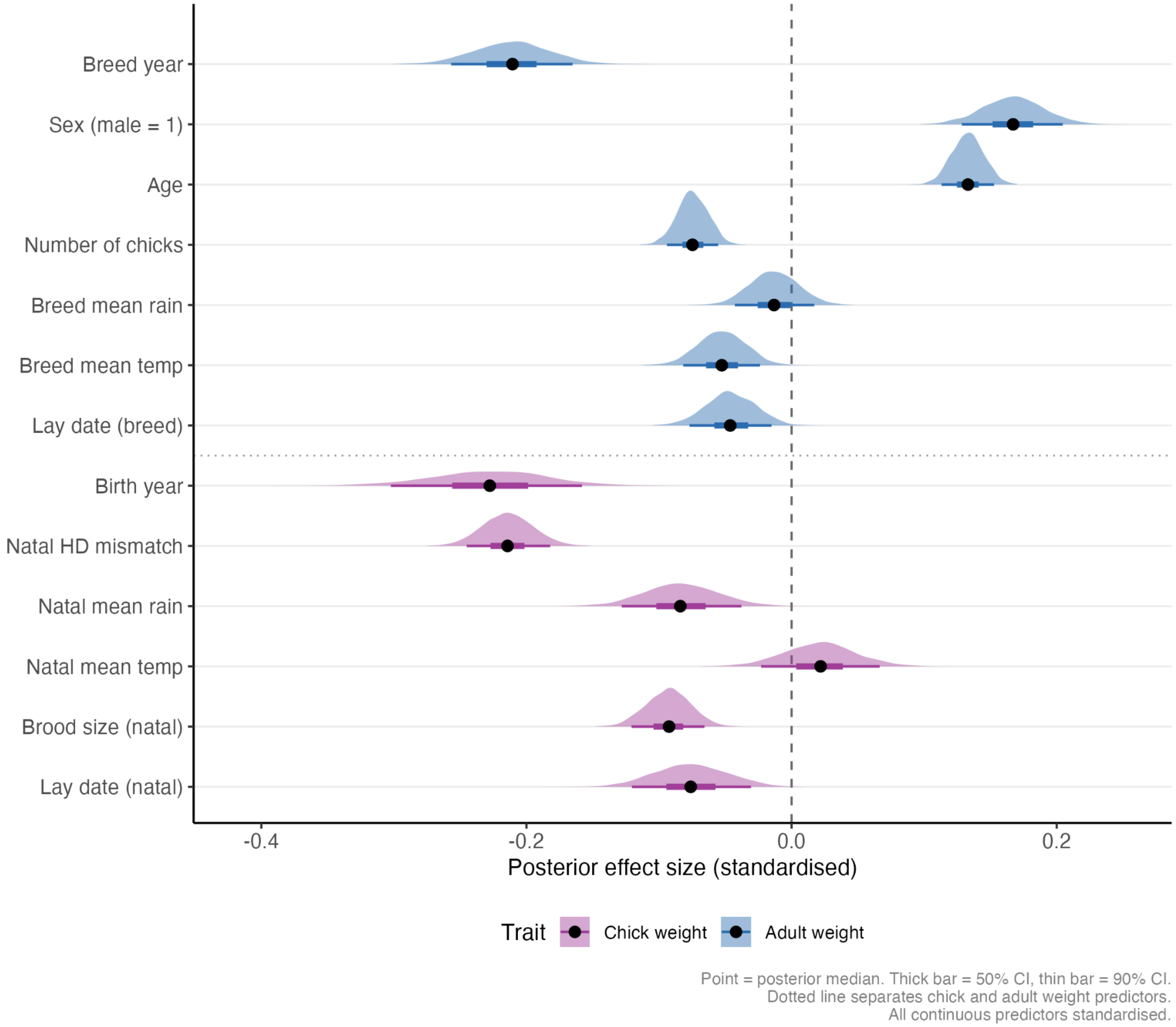


**Figure S10.** Posterior densities for fixed effects of standardised predictors on nestling mass (purple) and adult mass (blue). Points = posterior median; thick bar = 50% CI; thin bar = 90% CI; dashed line = zero).

### S5.2 Part 3 Genetic CRN models

#### Fixed effect posteriors

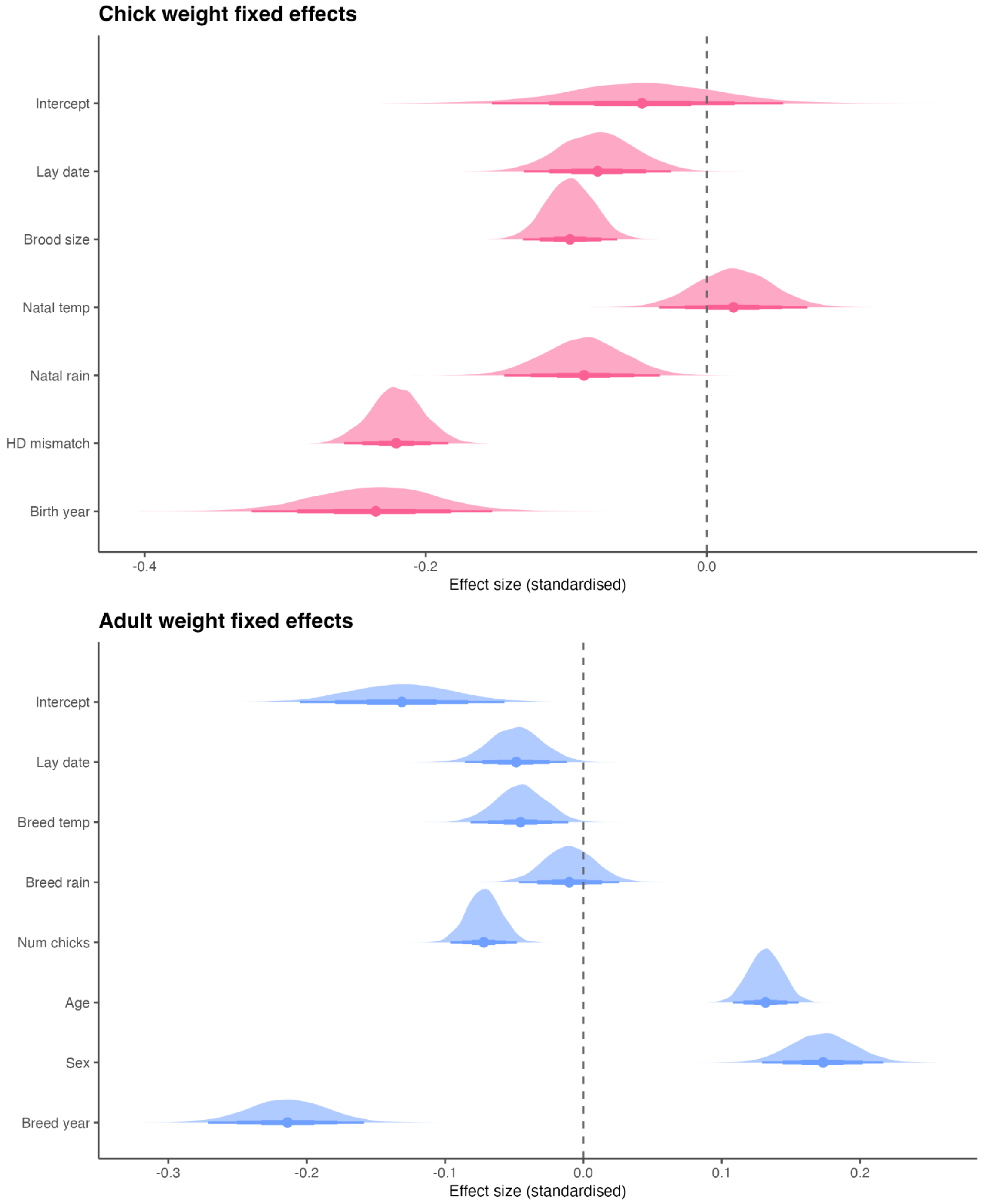


**Figure S11.** Posterior densities for fixed effects of standardised predictors on nestling mass (top) and adult mass (bottom). Points = posterior median; thick bar = 50% CI; thin bar = 90% CI; dashed line = zero).

### S6. Posterior predictive checks

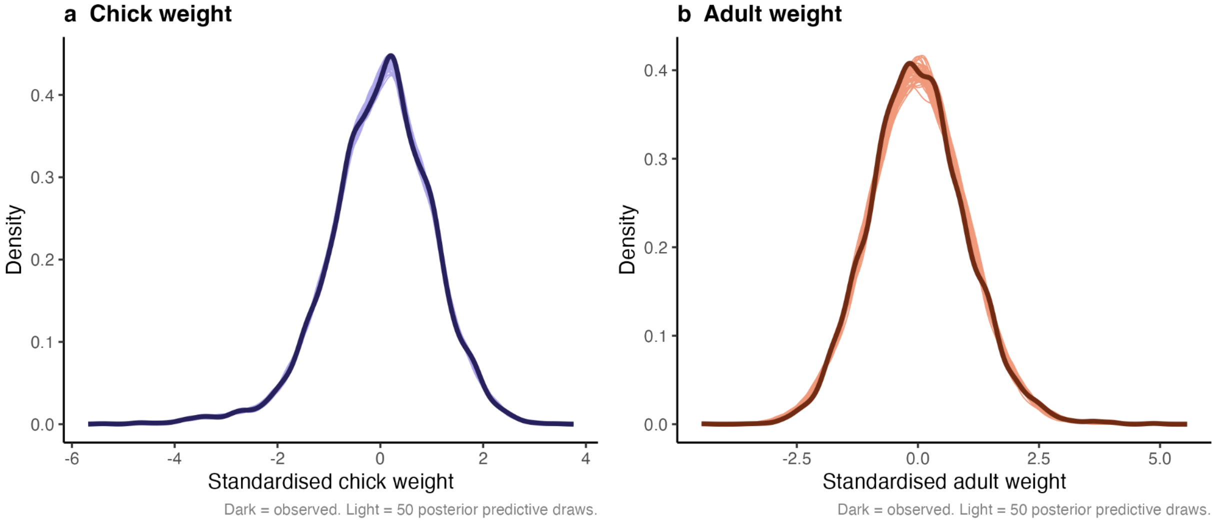


**Figure S12.** Posterior predictive checks showing the distributions of observed data (dark line) and data generated under the statistical model (faint lines), for a) nestling mass, and b) adult mass.

### S7. Genetic CRN model using quadratic terms for temperature

To test whether the null GxE result for $r_A$ reported in the main text is robust to the inclusion of quadratic terms as in Part 1 and 2, we parameterised the context matrix to have the following terms: intercept, natal_temp, adult_temp, natal_temp², adult_temp². Figures S13 and S14 show that the $r_E$ slope is credibly positive with breeding temperature, mirroring the phenotypic correlation identified in Part 1, and the $r_A$ slopes are all unchanged across the temperature gradients, with wide credible intervals overlapping zero.

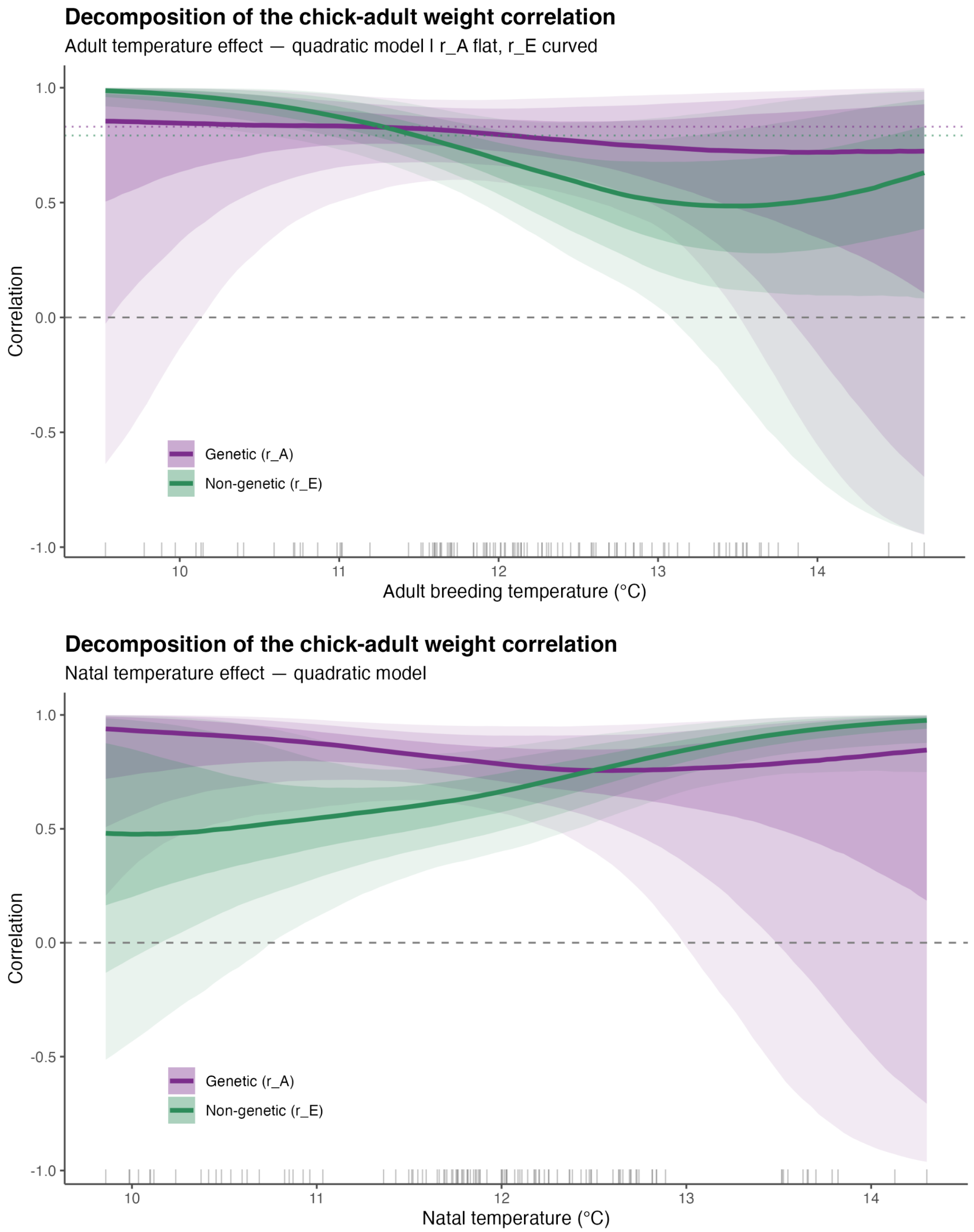


**Figure S13.** Reaction norms for the additive genetic correlation ($r_A$ in purple) and non-genetic among-individual correlation ($r_E$ in green) between nestling mass and adult body mass of great tits as functions of (A) adult breeding temperature, and (B) natal temperature (linear + quadratic terms).

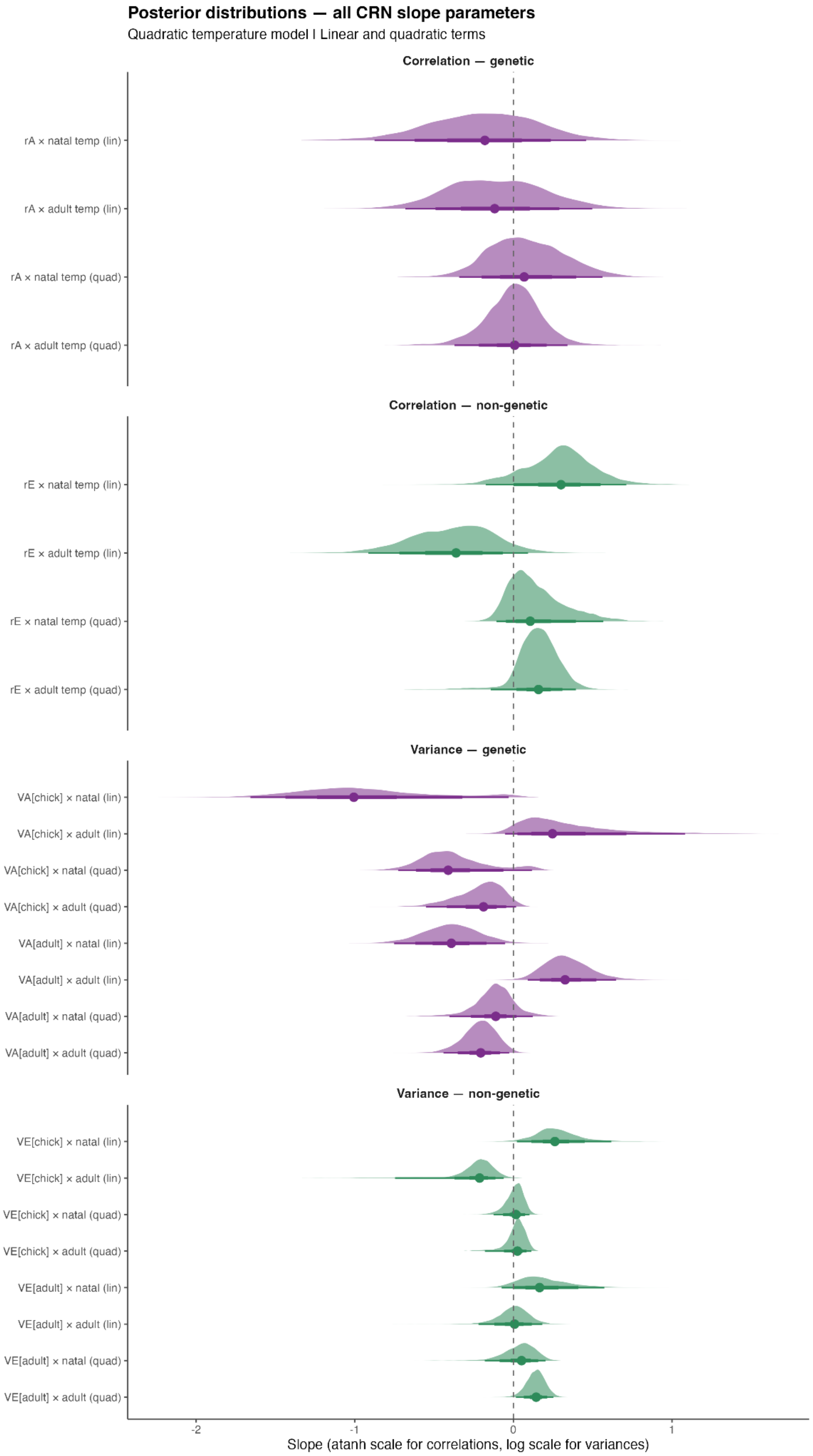


**Figure S14.** Posterior distributions of CRN and VRN slopes on the *atanh* and *log* scale respectively (point = posterior median; thick bar = 50% CI; thin bar = 90% CI; dashed line = zero).
